# Differential impact of FLASH and conventional radiotherapy on a pivotal metabolic organ: White Adipose Tissue

**DOI:** 10.64898/2026.03.30.715260

**Authors:** Gaia Scabia, Giulia Furini, Alice Usai, Giulia Asero, Emanuela Guerra, Eduarda Mota Da Silva, Claudia Kusmic, Andrea Cavalieri, Damiano Del Sarto, Mario Costa, Martin Wabitsch, Filippo Rossi, Roberta Di Pietro, Stefano Lattanzio, Tonia Luca, Salvatore Pezzino, Sergio Castorina, Roberto Cusano, Simone Capaccioli, Alessandra Gonnelli, Fabiola Paiar, Fabio Di Martino, Saverio Cinti, Margherita Maffei

## Abstract

**BACKGROUND:** Subcutaneous white adipose tissue (scWAT), a key metabolic and endocrine organ, is inevitably exposed during radiotherapy (RT). While RT is a cornerstone of cancer treatment, its efficacy is limited by toxicity to surrounding healthy tissues. Ultra-high dose rate (FLASH) RT has emerged as a promising modality capable of preserving tumor control while reducing normal tissue damage - the so-called FLASH effect. Clinical evidence indicates that childhood exposure to conventional (CONV) RT is associated with long-term dysmetabolism and WAT dysfunction. However, the impact of FLASH-RT on WAT has not been investigated.

**AIM:** To compare the effects of FLASH- and CONV-RT on adipocyte function and scWAT homeostasis, and to identify molecular and structural changes associated with each modality.

**METHODS:** We evaluated the effects of FLASH- and CONV-RT on adipocytes and scWAT using a dedicated linear accelerator capable of delivering both modalities. Experiments were performed in the human SGBS preadipocyte/adipocyte cell line and in a mouse model subjected to proximal hind limb irradiation, with analyses conducted 70 days post-exposure.

**RESULTS:** RT impaired adipogenic differentiation in a dose-dependent manner, with a relative sparing effect of FLASH at 4-8 Gy. Mature adipocytes exhibited radioresistance, with protection by FLASH at 8 Gy. In vivo, both regimens reduced fat mass without affecting body weight, with greater loss following CONV-RT. Transcriptomic profiling of scWAT revealed inflammatory and neurodegenerative signatures after CONV-RT, whereas FLASH-RT induced minimal transcriptional changes. Histological and ultrastructural analyses confirmed increased cellular damage, vacuolization, lipid spill-over, and reduced PLIN1 expression, predominantly in CONV-treated mice.

**CONCLUSIONS:** WAT homeostasis is sensitive to conventional RT, whereas FLASH-RT better preserves tissue structure and function, with implications for long-term metabolic health in cancer survivors.

## 1. Introduction

About half of cancer patients receive radiotherapy (RT) during their treatment. Despite its effectiveness in nearly 40% of cases, the dose and fractionation are constrained by collateral damage to healthy tissues. A transformative strategy is represented by ultra-high dose rate RT—known as FLASH—which preserves therapeutic efficacy in tumor control, while reducing toxicity to normal tissues, a phenomenon referred to as the FLASH effect [1–3]

Despite its clinical relevance, the impact of RT on white adipose tissue (WAT) remains poorly investigated. Like skin, WAT is invariably traversed by ionizing radiation (IR) and is widely distributed as a hypodermal layer. Importantly, WAT is recognized as the largest endocrine organ in the body, and alterations at this level are expected to exert systemic consequences with effects on multiple target organs [4–7]. In the context of cancer and RT, WAT plays an important role in modulating the organismal response. For instance, the adipokine adiponectin, secreted by adipocytes, has been shown to protect fibroblasts by attenuating radiation-induced cell death and senescence, while exerting no pro-proliferative effects on cancer cells [8]. Moreover, adipocytes are integral components of the tumor microenvironment and actively interact with malignant cells. Through the release of hormones, adipokines, growth factors, and proteases, they promote cancer invasion, angiogenesis, and the recruitment of inflammatory and vascular cells. Additionally, by modulating immune cell activity and interacting with fibroblasts and macrophages, adipocytes contribute to tumor-associated inflammation and extracellular matrix remodeling [9]. Clinical data also suggest that radiation to adipose-rich regions is detrimental, leading to an increased risk of cancer recurrence and metastasis compared to non-radiation interventions [10–12].

Further understanding of how RT impacts adipocyte function is therefore highly relevant—and indeed essential—for predicting its long-term effects and therapeutic efficacy. Mechanistic investigations on human preadipocytes have shown that exposure to medium doses of X-rays (10 Gy) negatively affects triglyceride accumulation during adipogenesis, although only minimal transcriptional changes were detected, with gene expression profiles largely resembling controls [13]. In rodent models, RT was found to reduce fat depot size and alter the expression of lipogenesis-associated genes [14,15]. Clinical evidence further supports a long-term impact of RT on WAT. Observational studies in adults who had received RT during childhood, prior to bone marrow transplantation, reported features of metabolic dysfunction, including insulin resistance, reduced adiponectin levels, increased visceral fat, dyslipidemia [16], and lipodystrophy [17]. More recently, Cohen and colleagues used bulk RNA-seq with single-cell–based deconvolution to uncover profound and persistent alterations in WAT of adults exposed to childhood RT, including features consistent with fibrosis, a relative reduction in progenitor cell fractions, and expansion of macrophage populations with a complex inflammatory and immunomodulatory phenotype [18].

Collectively, these findings indicate that RT induces irreversible alterations in WAT, underscoring the need for further mechanistic studies to elucidate how conventional (CONV) and FLASH-RT differentially affect this critical metabolic organ.

In this study, we sought to address this gap by systematically comparing the effects of CONV-and FLASH-RT on WAT morphology, differentiation capacity, and gene expression, both *in vitro* and in a murine model. Our data demonstrate that the sparing effect of FLASH-RT previously reported for skin [1,19] and other normal tissues [20–24] also extends to WAT. Moreover, we provide an in-depth characterization of the impact of ionizing radiation on WAT biology, contributing to a field of investigation that remains relatively unexplored.

## 2. Methods

### 2.1 Experimental Design

This study was designed to compare the effects of CONV electron RT and FLASH-RT on Simpson–Golabi–Behemel syndrome (SGBS) cells, a human preadipocyte model capable of differentiating into mature adipocytes upon exposure to a defined hormonal cocktail [25,26], irradiated at either the preadipocyte or adipocyte stage, as well as on WAT from C57BL/6J mice. Radiation was delivered using a novel linear accelerator (LINAC) specifically developed to ensure reproducible and finely tunable control of electron beam parameters. The dose distribution was relatively uniform within the first 1–2 cm from the surface, enabling evaluation of the biological effects of both CONV-RT and FLASH-RT in tissues located within this depth range, including subcutaneous (sc) WAT.

SGBS cells were used to assess the response to both irradiation modalities at 4, 8, and 16 Gy, when irradiated at either the preadipocyte or differentiated adipocyte stage (Figure 1A, B). Endpoints included cell survival, senescence, and adipogenic potential.

**Figure 1.**
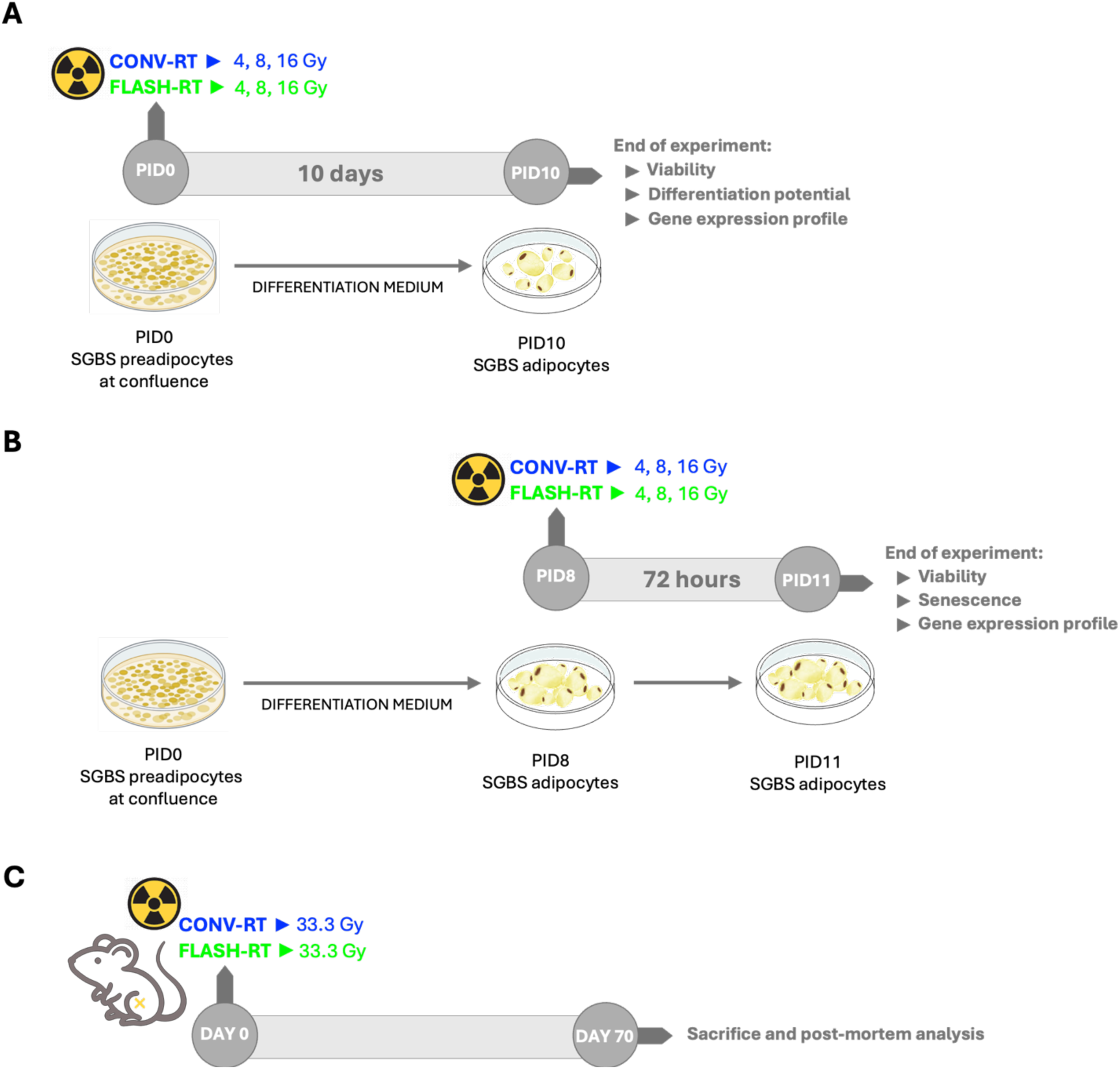
Experimental design schemes. **(A)** SGBS preadipocytes at confluence were either irradiated or non-irradiated with CONV- or FLASH-RT at three doses (4, 8 and 16 Gy), after which they were differentiated and analyzed after 10 days. **(B)** SGBS mature adipocytes were either irradiated or non-irradiated with CONV- or FLASH-RT at three doses (4, 8 and 16 Gy). Analysis was performed 72 hours thereafter. **(C)** Healthy C57Bl/6 mice were either non-irradiated or irradiated with CONV- or FLASH-RT (33.3 Gy) at the proximal left hindlimb to target the iscWAT, and were monitored for 70 days after RT.

Inguinal scWAT (iscWAT) was collected from C57BL/6J mice following irradiation with CONV-RT or FLASH-RT to assess the response to both modalities. Animals were randomly assigned to untreated (control, CNT), CONV, or FLASH groups (n = 3–5 mice per group). Mice in the CONV and FLASH groups received a single 33.3 Gy dose to the proximal left hind limb 1–2 days after housing (Day 0) and were sacrificed 70 days post-irradiation to evaluate mid- and long-term effects (Figure 1C). Animals reaching predefined humane endpoints due to health deterioration were euthanized earlier. The impact of CONV- and FLASH-RT on iscWAT was assessed by high-throughput gene expression profiling and histological evaluation using light and transmission electron microscopy (TEM). All data analyses were performed in a blinded manner.

### 2.2 Cells

SGBS human preadipocytes were cultured in DMEM-Ham’s F12 (1:1) medium containing 2 mM L-glutamine, 100 I.U./mL penicillin and 100 μg/mL streptomycin, 17 µM pantothenic acid, 33 µM biotin, and 10% of fetal bovine serum (FBS). Cells were seeded in 12-well or 96-well plates, depending on the analysis to be performed, using the appropriate volume of culture medium, and maintained at 37°C in a humidified atmosphere of 5% CO₂.

When complete confluence was reached (Post-Induction Day 0, PID0), white adipogenic differentiation was induced using a serum-free medium supplemented with 100 I.U./mL penicillin and 100 μg/mL streptomycin, 33 μM biotin, 17 μM pantothenic acid, 2 μM rosiglitazone, 10 μg/mL human apo-transferrin, 20 nM human insulin, 25 nM dexamethasone, 500 μM 3-isobutyl-1-methylxantine (IBMX), 100 nM cortisol, and 200 pM triiodothyronine. After four days (PID4), the medium was changed and rosiglitazone, dexamethasone, and IBMX were removed. Cells were maintained in the new medium (maintenance medium) for the remaining days of differentiation. The maintenance medium was replaced every third day.

To assess the effects of RT on adipogenic potential, SGBS preadipocytes were exposed to either CONV- or FLASH-RT at doses of 4, 8, or 16 Gy at PID0, followed by induction of differentiation. Cells were subsequently analyzed at PID10. To evaluate the impact of radiotherapy on mature adipocytes, differentiated SGBS adipocytes at PID8 were irradiated with CONV- or FLASH-RT at 4, 8, or 16 Gy and analyzed 72 hours post-irradiation (PID11).

### 2.3 Animals

Male C57BL/6J mice (6–7 weeks old) were purchased from Charles River Laboratories Italia SRL. Following a 2-week acclimatization period in the animal facility, animals were housed in groups of 3–4 per cage and then were randomly assigned to CNT, FLASH- and CONV-RT (N=3-5 mice per group) groups. Food and water were available *ad libitum*. Mice in CONV and FLASH groups received electron radiation with either modality (33.3 Gy total dose, DAY0) to the left leg (thigh area). 70 days after RT, mice were sacrificed. At the time of sacrifice, mice body weight was recorded, and fat depots (inguinal, epididymal and perirenal adipose tissues) were dissected and weighted. Inguinal fat depots (iscWAT) were then either snap-frozen in liquid nitrogen for RNA extraction or formalin-fixed for histological analysis. During all experimental period mice were monitored for their general health condition.

### 2.4 Irradiation Study

All the irradiations were performed at the Centro Pisano FLASH Radiotherapy (CPFR) using the ElectronFlash linear accelerator [27], which delivers electron beams of 7 or 9 MeV with the possibility of modifying the average dose-rate (ADR) and dose per pulse (DPP) by varying the e-beam current and the pulse repetition frequency (PRF). This setup enables switching the irradiation conditions from conventional (CONV) to FLASH while maintaining the same experimental configuration. All irradiation parameters are summarized in Tables 1 and Table 2 for the in vitro and in vivo studies, respectively.

**Table 1.** In vitro radiotherapy experimental parameters: relevant parameters used in each irradiation condition.

|  | Number of pulses | DPP [Gy] | Total dose [Gy] | Pulse duration [µs] | PRF [Hz] | ADR [Gy/s] |
| --- | --- | --- | --- | --- | --- | --- |
| **FLASH-RT** | 4, 8, 16 | 1 | 4, 8, 16 | 4 | 180, 210, 225 | 240 |
| **CONV-RT** | 182, 364, 728 | 0,022 | 4, 8, 16 | 4 | 3 | 0,06 |

*DPP, dose per pulse; PRF, pulse repetition frequency; ADR, average dose rate*

**Table 2.** In vivo radiotherapy experimental parameters: relevant parameters used in each irradiation condition.

| | Number of pulses | DPP [Gy] | Total dose [Gy] | Pulse duration [μs] | PRF [Hz] | ADR [Gy/s] |
| --- | --- | --- | --- | --- | --- | --- |
| **FLASH-RT** | 7 | 4.75 | 33.3 | 4 | 170 | 940 |
| **CONV-RT** | 180 | 0.19 | 33.3 | 4 | 1 | 0.18 |

*DPP, dose per pulse; PRF, pulse repetition frequency, ADR, average dose rate*

For the *in vivo* study, all irradiations were performed on anesthetized mice (ketamine 100 mg/kg / xylazine 10 mg/kg, i.p.) using a collimator with 10 mm diameter centered on the animal’s left thigh. This choice allows the precise positioning of the electron beam on the region of interest (7 mm diameter). As described elsewhere [19], the animal was fixed by its teeth on a 3D printed support perpendicular to the applicator, so that the positioning of each animal would be reproducible.

The dosimetric characterization of our beams, carried out using the fD detector *[28]* and other measurement methods developed at CPFR and/or provided by other centers *[29–37]*, has been described in detail in our previous work [19].

### 2.5 Cell viability

Cell viability of irradiated and non-irradiated (CNT) SGBS cells was assessed at the mature adipocyte stage in cells irradiated at PID0 (preadipocyte stage) and PID8 (mature adipocyte stage), as described in the Experimental design section. Briefly, cells were incubated with 2.5 µM propidium iodide for 30 minutes at 37 °C, followed by fixation with 4% cold paraformaldehyde for 15 minutes at room temperature (RT). After 3 washes with Dulbecco’s phosphate-buffered saline (DPBS), nuclei were counterstained with DAPI (#D1306, Thermo Fisher Scientific) according to the manufacturer’s instructions and imaged using a Nikon Eclipse Ti2 fluorescent microscope. Viable cells were identified as nuclei negative for propidium iodide staining and quantified using ImageJ software.

### 2.6 Oil Red O staining

The intracellular lipid content of irradiated and non-irradiated (CNT) SGBS cells was assessed by Oil Red O (ORO) staining. Cells were washed with DPBS, fixed with 10% formalin in DPBS for 1 hour at RT, and subsequently washed twice with Milli-Q water and once with 60% isopropanol for 5 minutes each. The ORO stock solution was prepared by dissolving 0.2 g of Oil Red O powder (#O0625, Sigma-Aldrich) in 45 mL of isopropanol at 30 °C, followed by cooling to RT and filtration through a 0.22 μm membrane. The working solution was prepared by mixing the stock solution with Milli-Q water in a 3:2 ratio. After fixation, isopropanol was removed and the ORO working solution was added to the cultures for 10 minutes, followed by five rinses with Milli-Q water. Brightfield images were acquired using a Leica DM IL LED inverted microscope with a Leica ICC50 HD camera. Lipid aggregates appeared as red droplets within the cytoplasm. After imaging, water was removed, and the plates were air-dried. ORO dye was eluted with 100% isopropanol for 10 minutes under gentle shaking, and the absorbance of the eluate was measured at 492 nm.

### 2.7 Senescence β-galactosidase cell staining

Senescence-associated β-galactosidase activity was evaluated cytochemically using the Senescence β-Galactosidase Staining Kit (#9860, Cell Signaling Technology), following the manufacturer’s instructions. Briefly, cells were rinsed with DPBS and fixed for 15 minutes. After fixation, the cells were washed twice with PBS. The β-galactosidase staining solution (pH 6.0) was then added to each well, and the cells were incubated overnight at 37 °C in a dry incubator. Stained cells were imaged using the optical microscope Leica DM IL LED, and the proportion of senescent cells was determined by quantifying β-galactosidase-positive cells.

### 2.8 RNA extraction and analysis in SGBS cells and mouse iscWAT

*SGBS cells* - At the end of experimental period, SGBS cells (3-4 wells for experimental condition in a 12-well plate) were detached and resuspended in TRIzol reagent (#15596-018, Thermo Fisher Scientific).

*iscWAT* - Homogenization of frozen mice iscWAT (10-15 mg) collected from the left inguinal region of CNT, CONV and FLASH-irradiated mice was performed with a Qiagen TissueLyser II in 700 µL of TRIzol reagent.

The following procedure was applied for both types of samples. After homogenization/resuspension in TRIzol reagent, samples were processed for the isolation of RNA. The aqueous phase obtained after chloroform addition and centrifugation (12000 × *g* for 15 min at 4 °C) was loaded onto Direct-zol RNA Microprep columns (# R2060, Zymo Research) and RNA eluted in 15 uL RNAse-free water. The RNA amount was quantified by Nanodrop (Qiagen). cDNA was synthesized from 0.2 μg of total RNA using the iScript™ Reverse Transcription Supermix for RT-qPCR (#1708840, Bio-Rad Laboratories). Droplet digital PCR (ddPCR) was subsequently performed. Briefly, PCR reaction mixtures were prepared for each sample using 10 ng of cDNA, the optimal amount of probes (defined in preliminary tests), and ddPCR™ Multiplex Supermix (#12005910, Bio-Rad Laboratories). The mixtures were loaded into disposable cartridges (#1864008, Bio-Rad Laboratories) together with 70 μl of Droplet Generation Oil (#1863005, Bio-Rad Laboratories) and droplets were generated using the QX200™ Droplet Generator (Bio-Rad Laboratories). A total of 40 μl of droplets per sample was transferred to a 96-well PCR plate for end-point amplification. Following PCR, the plate was loaded into the QX200™ Droplet Reader (Bio-Rad Laboratories) for detection and quantification of positive droplets.

Absolute quantification of target genes and the housekeeping gene *TATA Binding Protein* (*TBP*) was performed using QuantaSoft™ Software v1.7 (Bio-Rad Laboratories). The following genes were analyzed:

- FAM-conjugated probes (Applied Biosystems, USA): *HP* (Hs 00978377_m1), *IL6* (Hs 00985639_m1), *MCP1* (Hs 00234140_m1), *FABP4* (Hs 01086177_m1), *ADIPOQ* (Hs 00605917_m1), *LEP* (Hs 00174877_m1), *TBP* (Hs 00427620_m1), *CDKN1A* (p21) (dHsaCPE5052298), *COL1A1*(Hs00164004_m1), *COL3A1*(Hs00943809_m1), *COL6A3* (Hs00915125_m1), *TGFB* (Hs00998133_m1).
- HEX-conjugated probes (Bio-Rad Laboratories, USA): *PPARG* (dHsaCPE5036471), *FASN* (dHsaCPE5047919), *PLIN1*(dHsaCPE5040023), *SOD* (dHsaCPE5029679), *NOX* (dHsaCPE5032151), *CDKN2A* (p16)(dHsaCPE5045105).

Gene expression levels were normalized to *Tbp* and expressed as fold change relative to control samples.

### 2.9 Transcriptomic analysis

Bulk RNA sequencing of irradiated and non-irradiated iscWAT was performed at the CRS4 facility. Total RNA was, extracted as described above, was assessed for quantity and quality (RIN ≥ 7.5) prior to library preparation. Ribosomal RNA was depleted, and strand-specific libraries were generated, individually indexed, and pooled for paired-end sequencing on an Illumina NovaSeq X Plus platform. Sequencing reads were processed using the in-house RiDE pipeline (https://github.com/solida-core/ride), developed at CRS4, which performs alignment, quantification, and quality control. Differential expression analysis was conducted using DESeq2 [38], followed by functional enrichment analysis with DAVID (https://david.ncifcrf.gov/). Detailed procedures for RNA-seq and bioinformatics analyses have been described previously [19]. RNA-seq datasets are available under accession number GSEXXXX in the Gene Expression Omnibus (NCBI).

### 2.10 Light microscopy

#### Sample preparation

iscWAT samples from left hindlimb of treated and CNT mice were processed for histological analysis as previously described in detail [39]. Briefly, tissues were fixed in 4% (w/v) paraformaldehyde in 0.1 M phosphate buffer at pH 7.4 for 24 h at 4°C. Samples were then dehydrated in increasing concentrations of alcohol, clarified in xylene and then embedded in paraffin blocks. From each sample, 3 μm-thick sections were cut using a sliding microtome, mounted onto glass slides, air-dried, and processed for further analyses.

#### Histochemistry with morphometry

Fibrosis quantification was performed using Sirius Red staining [40]. Tissue sections were dewaxed in xylene and rehydrated through sequential immersions in alcohol solutions of decreasing concentration. The sections were then incubated with Picro-Sirius Red dye for one hour at room temperature and then abundantly washed and contrasted with hematoxylin for 2 min. Samples were then rinsed, dehydrated, and mounted using Eukitt® Mounting Medium (#03989, Merk Life Science S.r.l). Images of Sirius Red-stained tissues were captured at 2.5x magnification, to acquire a number of images corresponding to the full size of the tissue section. Semi-automated image analysis was performed to quantify fibrosis, using ImageJ software. Images were converted to grayscale, and red-stained areas (indicating presence of fibrosis) were selected using the threshold tool. Fibrosis was expressed as the percentage of Sirius red-positive area relative to the total fat tissue area.

#### Immunohistochemistry with morphometry

Tissue sections (3 μm thick) were processed by BenchMark ULTRA system (Roche Tissue Diagnostics, Italy), following manufacturer’s instructions. The fully automated system with individual slide drawers allows to stain each single slide. The overall workflow was performed using Ventana Ultra CC1 buffer (Roche Tissue Diagnostics), followed by automated addition of ready-to-use CONFIRM anti-CD68 (KP-1) Rabbit Monoclonal Primary Antibody (Roche Tissue Diagnostics) or manual addition of rabbit anti-perilipin1 (ab3526, 1:400 in PBS; Abcam). The UltraView Universal DAB Detection Kit (Roche Tissue Diagnostics) was then applied automatically, followed by hematoxylin counterstaining for nuclei and mounting. Tissue sections were examined under a light microscope (Zeiss Axioskop 40; Carl Zeiss GmbH), and digital images were acquired using a Zeiss Axiocam 503 color camera and used for morphometric analyses.

The evaluation of CD68-positive cells was performed acquiring 20 random fields per tissue section at 40x magnification. CD68-positive macrophages were then counted using ImageJ software and normalized to 10^4^ adipocytes.

For the determination of adipocyte viability [41–43], sections were captured at 2.5× magnification to acquire images covering the full size of each tissue section. ImageJ software was used to calculate the total adipose tissue area, composed of PLIN1-positive and PLIN1-negative adipocytes. The percentage of PLIN1-negative adipocytes was then calculated for each sample.

### 2.10 Transmission Electron Microscopy (TEM)

Murine iscWAT samples (∼1 mm²) were fixed in 2.5% glutaraldehyde and processed for high-resolution/high-magnification microscopy. Semithin (800 nm) and ultrathin (∼80 nm) slices were obtained from resin-embedded tissues using a PowerX ultramicrotome (RMC Products). Semithin sections were stained with 1% toluidine blue and 1% sodium tetraborate. Whole sections were digitalized using a NanoZoomer-SQ digital slide scanner (Hamamatsu Photonics). Ultrathin sections were mounted on 200 mesh copper grids and counterstained with UranyLess and lead citrate. TEM analysis was performed using a JEOL JAM-1400 Flash microscope operated at 120 kV. Morphometric analyses of adipocytes size and vacuolization were performed on the whole digitalized toluidine-blue stained semithin sections using the Hamamatsu NDP view 2.0 image analysis software.

### 2.11 Statistical analyses

Graphical representation and statistical analyses were performed using GraphPad Prism. Data are presented as mean±SEM. All statistical analyses were performed using GraphPad Prism. Normality was assessed using the Shapiro–Wilk test, and homogeneity of variances using the Brown–Forsythe test. Depending on the experimental design, comparisons were performed using one-way ANOVA for single-factor analyses (or the non-parametric Kruskal–Wallis test when ANOVA assumptions were violated) or two-way ANOVA for experiments involving two factors. For experiments in which multiple cells were analyzed per subject, a nested ANOVA (cells within subjects) was performed. Multiple comparisons between groups were adjusted using Tukey’s post hoc test.

Correlations between variables were evaluated using Pearson’s correlation coefficient (r), with associated p values reported. All tests were two-tailed, and p values < 0.05 were considered statistically significant.

## 3. Results

### 3.1 Impact of RT on Adipocyte Survival and Adipogenesis: Comparison Between CONV-and FLASH-RT

We initially investigated the effects of CONV- and FLASH-RT *in vitro*, using the human SGBS cell line, both in its undifferentiated (PID0) and differentiated state (PID10). To evaluate the impact of RT on adipogenic potential, PID0 SGBS cells were exposed to 3 radiation doses (4, 8, and 16 Gy). Adipogenic differentiation was assessed 10 days later (PID10) and quantified by ORO staining to detect intracellular triglyceride (TG) accumulation. Importantly, irradiation did not compromise cell viability: no significant differences in total nuclei counts at PID10 were observed between irradiated and non-irradiated cells, irrespective of the RT modality (Figure 2A). In contrast, TG accumulation was clearly reduced by RT, with the strongest effect observed at the highest dose (16 Gy) for both modalities (Figure 2B). Notably, at low-to-intermediate doses (4 and 8 Gy), the differentiation potential was significantly better preserved following FLASH-RT compared to CONV-RT.

**Figure 2.**
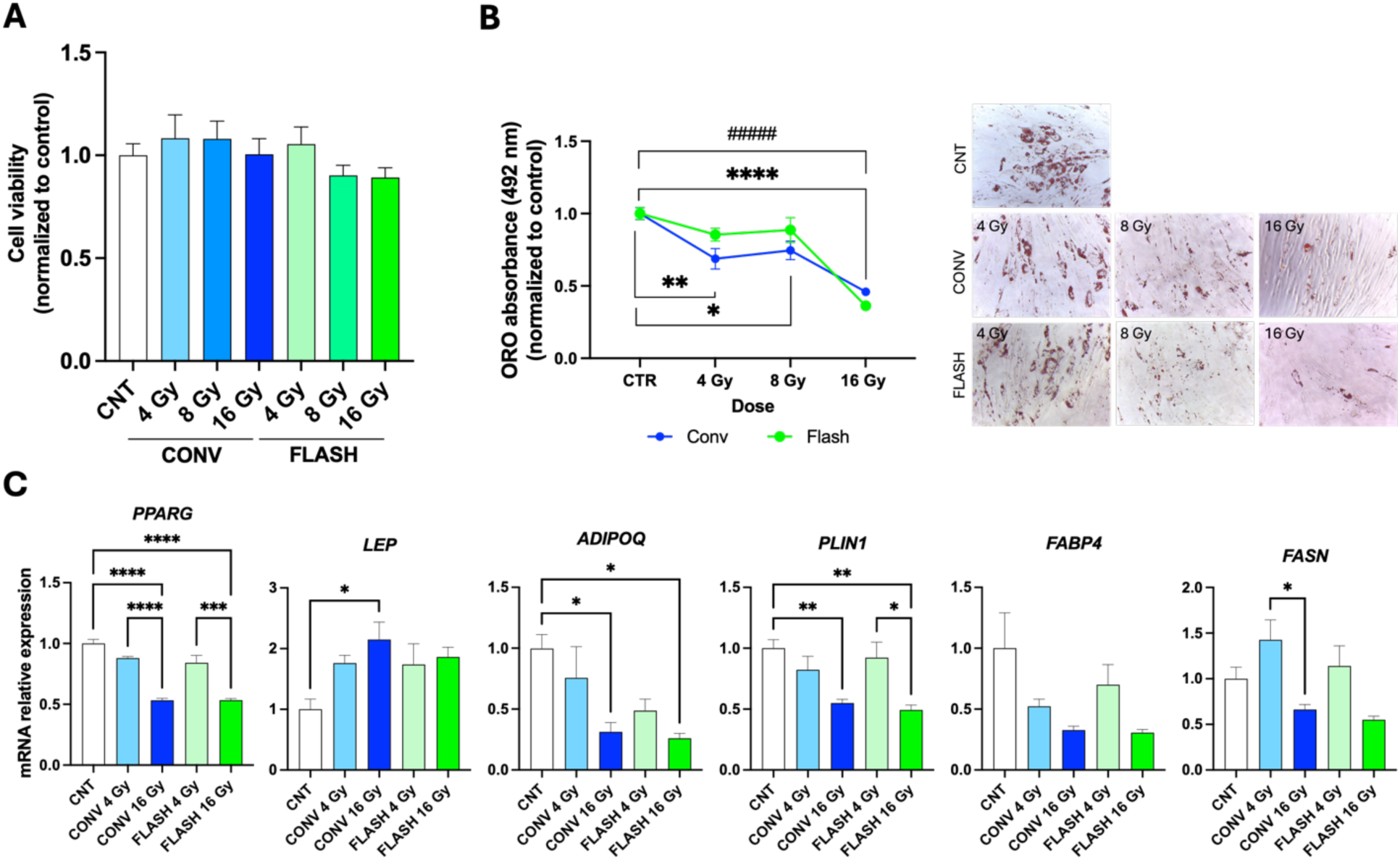
CONV- and FLASH-RT effect on SGBS preadipocytes and on adipogenesis. **(A)** Cell viability of PID10 SGBS cells in non-irradiated controls (CNT) and in CONV- or FLASH-irradiated cells (4, 8, and 16 Gy). A minimum of nine fields were analyzed for each experimental condition. **(B)** Left: Oil Red O (ORO) absorbance in CNT and CONV- or FLASH-irradiated (4, 8, and 16 Gy) PID10 SGBS cells, used to quantify triglyceride (TG) accumulation efficiency (* = CNT vs CONV; # = CNT vs FLASH). A minimum of four wells were analyzed for each experimental condition. Right: Representative images of ORO-stained PID10 SGBS cells under all experimental conditions. **(C)** Gene expression analysis of markers associated with mature adipocytes in CNT and CONV- or FLASH-irradiated (4 and 16 Gy) PID10 SGBS cells. mRNA counts as obtained by ddPCR are normalized to CNT values. Data are presented as mean ± SEM. Statistical analysis was performed using two-way ANOVA, followed by multiple-comparison tests. *p < 0.05; **p < 0.01; ***p < 0.001; ****/#### p < 0.0001.

To gain further evidence of radiation-induced changes in adipogenesis, we monitored the expression of markers typically associated with mature adipocytes. These included the adipokines *LEP* and *ADIPOQ*, the key regulator of adipogenesis *PPARG*, and genes involved in TG accumulation and lipid droplet homeostasis—namely *FABP4*, *FASN*, and *PLIN1*.

Gene expression was assessed for samples irradiated with 4 Gy, where a difference in TG accumulation between CONV- and FLASH-RT was observed, and 16 Gy, where both modalities markedly suppressed TG accumulation. At 16 Gy, this effect was partly reflected at the transcriptional level, with a significant downregulation of some terminal adipocyte differentiation markers, regardless of radiation modality. A similar, albeit more moderate, effect was observed at 4 Gy; however, no clear differences between CONV- and FLASH-RT emerged (Figure 2C).

We next wanted to assess the effect of CONV- and FLASH-RT administered to mature adipocytes. In this case the irradiation at 3 different doses (4, 8 and 16 Gy) was performed at PID8, when SGBS had reached terminal differentiation. Cells were analyzed 72 hours thereafter (PID11) and monitored for cell death, senescence and gene expression. Our data demonstrate a substantial radio-resistance of these cells under both irradiation modalities with no significant difference in viability at any dose compared to CNT. Moderately higher adipocyte viability was observed for FLASH-RT compared with CONV-RT at 8 Gy (Figure 3A). We next investigated potential cellular damage by assessing senescence, a process known to drive multiple WAT dysfunctions, including impaired adipogenesis, chronic inflammation, aberrant adipocytokine production, and insulin resistance [44]. The irradiated groups displayed increased β-galactosidase staining—a canonical marker of senescence—with a more pronounced effect in CONV compared to FLASH-irradiated samples (Figure 3B). Consistently, quantitative PCR revealed upregulation of the acute stress response senescent marker p21 in cells irradiated at 8 Gy, regardless of modality. No changes were detected in p16: this is somewhat expected as this cyclin-dependent kinase inhibitor does not exert a major role in adipocytes [45] (Figure 3C). Analysis of inflammatory markers showed a significant upregulation of *MCP1* in both CONV- and FLASH-RT samples compared to CNT, with no changes in *IL6* and a moderate, albeit not significant, increase in *HP* (Figure 3D). Genes involved in adipocyte function (*LEP*, *ADIPOQ*, *PPARG*, *FABP4*, *FASN*, *PLIN1*), fibrosis (*COL1A1*, *COL3A1*, *COL6A3*, *TGFB1*), and oxidative stress response (*NOX*, *SOD*) showed no significant alterations following RT (Suppl. Figure 1).

**Figure 3.**
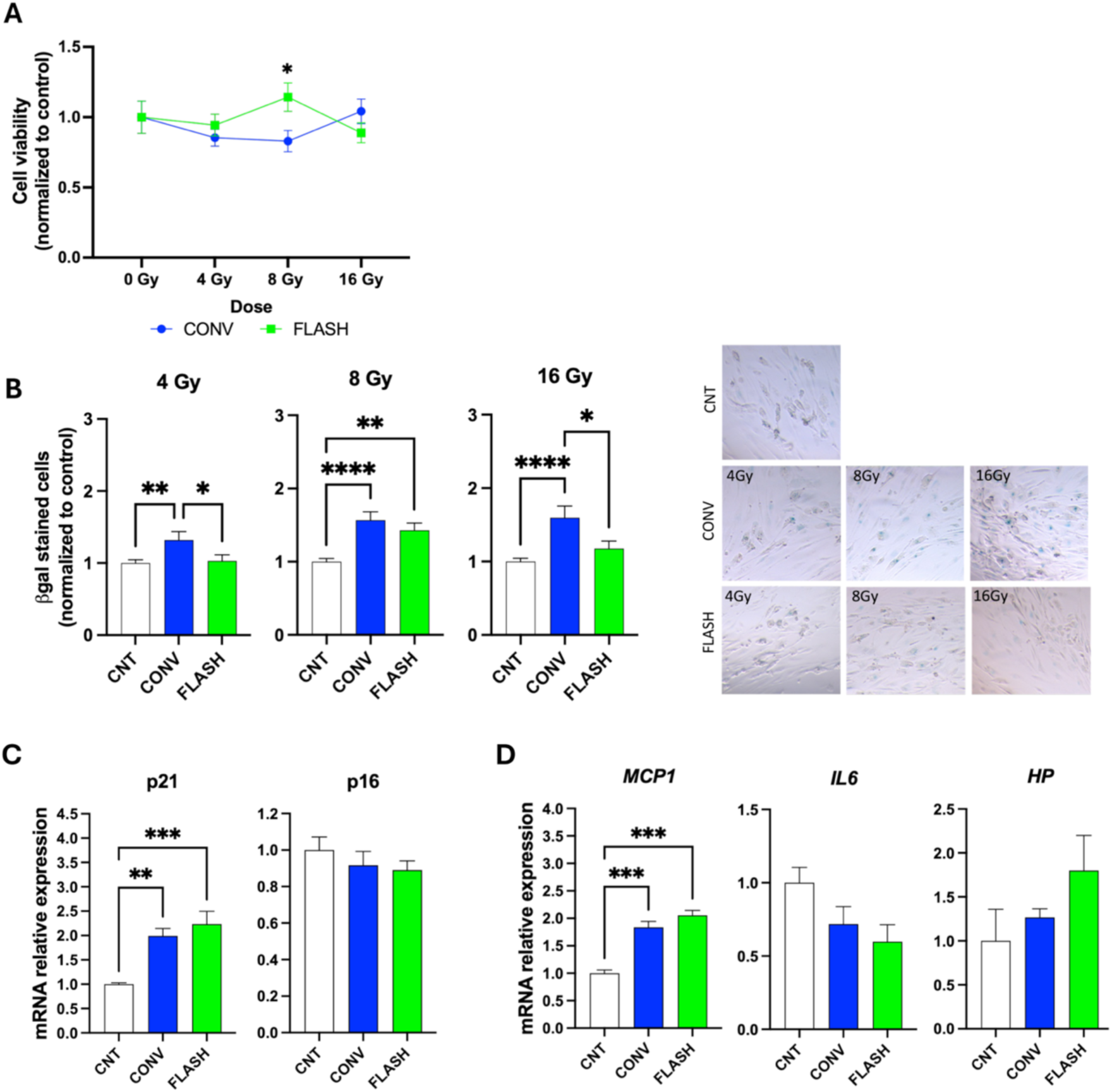
CONV- and FLASH-RT impact on adipocyte survival and senescence. **(A)** Viability of non-irradiated (CNT) and CONV- or FLASH-irradiated PID10 SGBS cells (4, 8, and 16 Gy). A minimum of nine fields were analyzed for each experimental condition. **(B)** Left: quantification of senescence in PID10 SGBS cells based on β-galactosidase staining at 4, 8, and 16 Gy. A minimum of nine fields were analyzed for each experimental condition. Right: representative images of PID10 SGBS cells under all experimental conditions, showing β-galactosidase-positive cells (blue). **(C–D)** Gene expression analysis of senescence markers (C) and inflammation-associated genes (D) in not-irradiated (CNT) and CONV- and FLASH-irradiated (8 Gy) PID10 SGBS cells. Data are presented as mean ± SEM. Statistical analysis was performed using two-way ANOVA, one-way ANOVA or Kruskal–Wallis test, as appropriate, followed by multiple-comparison tests. *p < 0.05; **p < 0.01; ***p < 0.001; **** p < 0.0001.

Collectively, these data indicate that *in vitro* RT impairs adipogenic potential and, to a lesser extent, activates pathways associated with cellular senescence. Notably, both effects appear to be attenuated under FLASH-RT conditions.

### 3.2 Impact of CONV and FLASH-RT on WAT in vivo

We next sought to evaluate how CONV- and FLASH-RT influence scWAT *in vivo*. To this end, we exposed mice to FLASH- or CONV-RT at a dose of 33.3 Gy, targeting the iscWAT as the primary tissue of interest (Figure 1C). Experimental details are provided in the Materials and Methods section and were described elsewhere [19]

#### 3.2.1 Size

At the end of the experimental period, the weight of iscWAT tended to decrease in irradiated mice despite overlapping values for body weight (BW) across the three experimental groups (Figure 4A and B). Interestingly, alterations were not limited to scWAT depots within the irradiated field, as a similar reduction was also detected in epididymal (EPI) and perirenal (PERI) WAT with both fat depots showing a more marked reduction following CONV-RT (Figure 4C). When the ratio between fat and body weight was calculated, differences became even more evident (Figure 4D and E). Consistent with the macroscopic findings, morphometric analysis of iscWAT adipocyte area on semithin sections revealed that adipocytes from CONV-RT mice had a smaller area compared with those from both control and FLASH-RT groups (Figure 4F).

**Figure 4.**
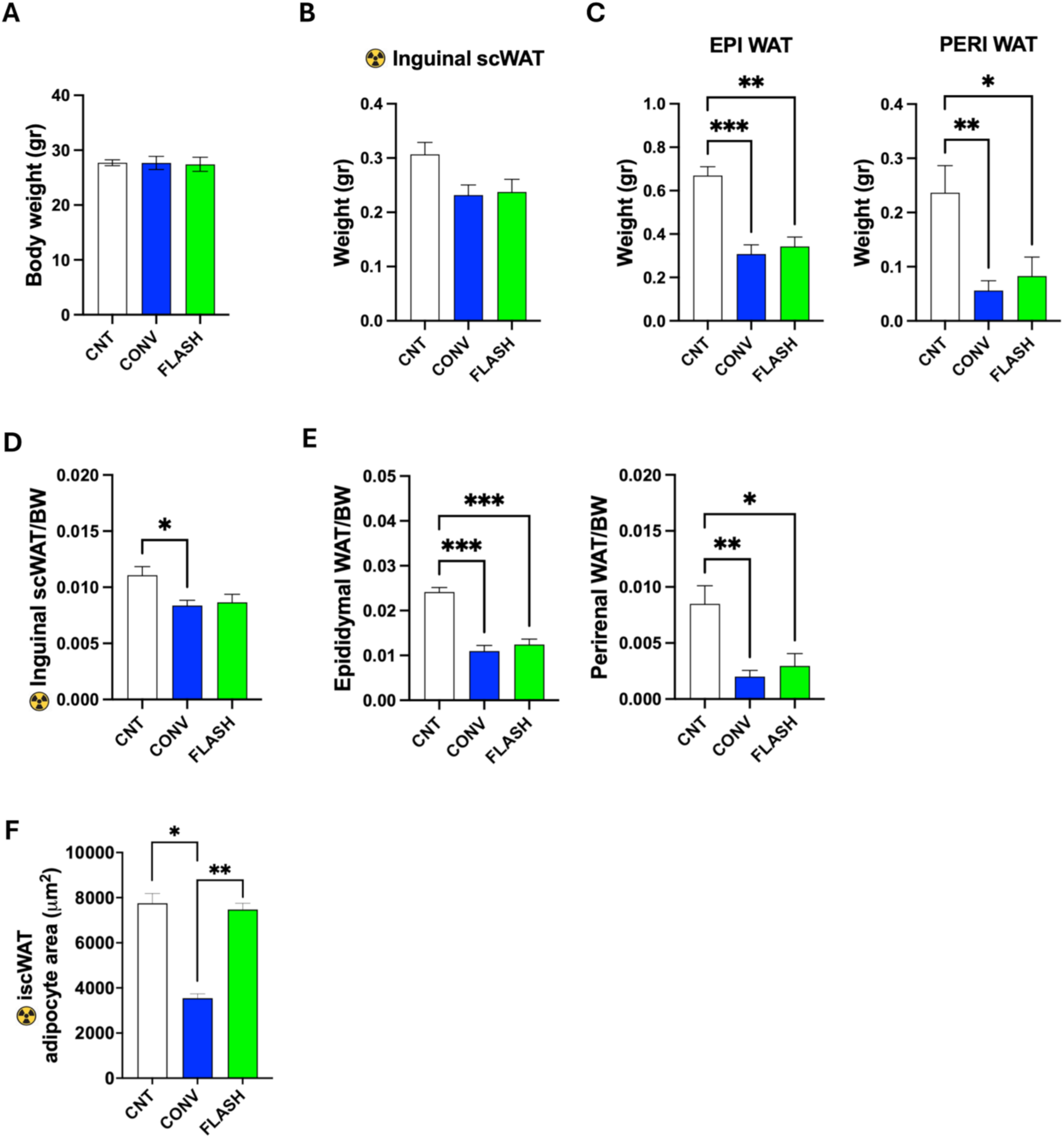
*In vivo* CONV- and FLASH-RT effect on white adipose tissue. **(A)** Body weight of CNT, CONV-RT and FLASH-RT mice at the end of the experimental period. **(B)** Weight of inguinal scWAT, the depot directly irradiated, of CNT, CONV-RT and FLASH-RT mice at the end of the experimental period. **(C)** EPI and PERI WAT weights of CNT, CONV-RT and FLASH-RT mice at the end of the experimental period. **(D-E)** Ratios of iscWAT depot weights (isc, EPI, and PERI) to BW at the end of the experimental period. **(F)** Morphometric analysis of iscWAT adipocytes area performed on the entire semithin section. Only well-preserved adipocytes were analyzed with NDPview_2 Software. Data are presented as mean ± SEM. Statistical analysis was performed using one-way ANOVA or nested one-way ANOVA, with subjects nested within experimental groups, followed by multiple-comparison test. *p < 0.05, **p < 0.01; ***p < 0.001. Data are presented as mean ± SEM.

#### 3.2.2 Gene expression profile

We next sought to evaluate if and how CONV- and FLASH-RT impacted on iscWAT transcriptome by performing bulk RNA sequencing (RNA-seq) on samples collected from irradiated hindlimbs 70 days after treatment. The analysis revealed a striking contrast in gene expression profiles between FLASH- and CONV-RT (Figure 5A, B; Supplementary Tables 1-3). Specifically, CONV-RT resulted in 172 differentially expressed genes compared to CNT, with 103 genes upregulated and 69 downregulated (Supplementary Tables 1, 2). In contrast, FLASH-RT induced differential expression of only 3 genes: Gm20708 (a predicted gene), upregulated, Vat1l (vesicle amine transport protein 1-like) and En1 (engrailed 1), both downregulated (Supplementary Table 3). Notably, En1 was the only gene commonly downregulated by both irradiation modalities (Figure 5B). En1 encodes a protein previously shown to be upregulated during brown adipogenesis, and promotes this process by enhancing the expression of key adipogenic and thermogenic regulators such as PPARγ2, C/EBPα, UCP1, and PGC1α [46]. Importantly, lipid accumulation in brown adipocytes is significantly reduced following EN1 knockdown [46]. Enrichment analysis identified 74 Gene Ontology (GO) Biological Process (BP) terms in the list of upregulated genes in CONV-RT, (p<0.05) (Figure 5C, Suppl. Figure 2). These were primarily related to inflammation and innate/adaptive immune responses, as well as chemotaxis and cell migration, consistent with chronic inflammation in irradiated iscWAT and immune cell recruitment, with enrichment of phagocytosis-related processes suggesting macrophage involvement. In addition, pathways related to oxidative stress and pro-inflammatory signaling were enriched, notably NF-κB, TNF, JNK-MAPK, JAK-STAT, and ERK1/2. In contrast, the 22 enriched GO BP terms among CONV-RT downregulated genes (Figure 5C) show a reduction in neuronal processes, as well as decreased angiogenesis and vascular regulation, pointing to a compromised neurovascular environment. Importantly, Smoothened (Smo) signalling pathway, a component of the broader Hedgehog signalling pathway involved in cellular differentiation, was enriched among the downregulated genes and specifically linked to Gli2 (GLI Family Zinc Finger 2), Cdon (Cell Adhesion Associated, Oncogene Regulated), and Tuba1a (Tubulin Alpha 1a). This enrichment may reflect a reduction in adipocyte differentiation signaling within adipose tissue [47].

**Figure 5.**
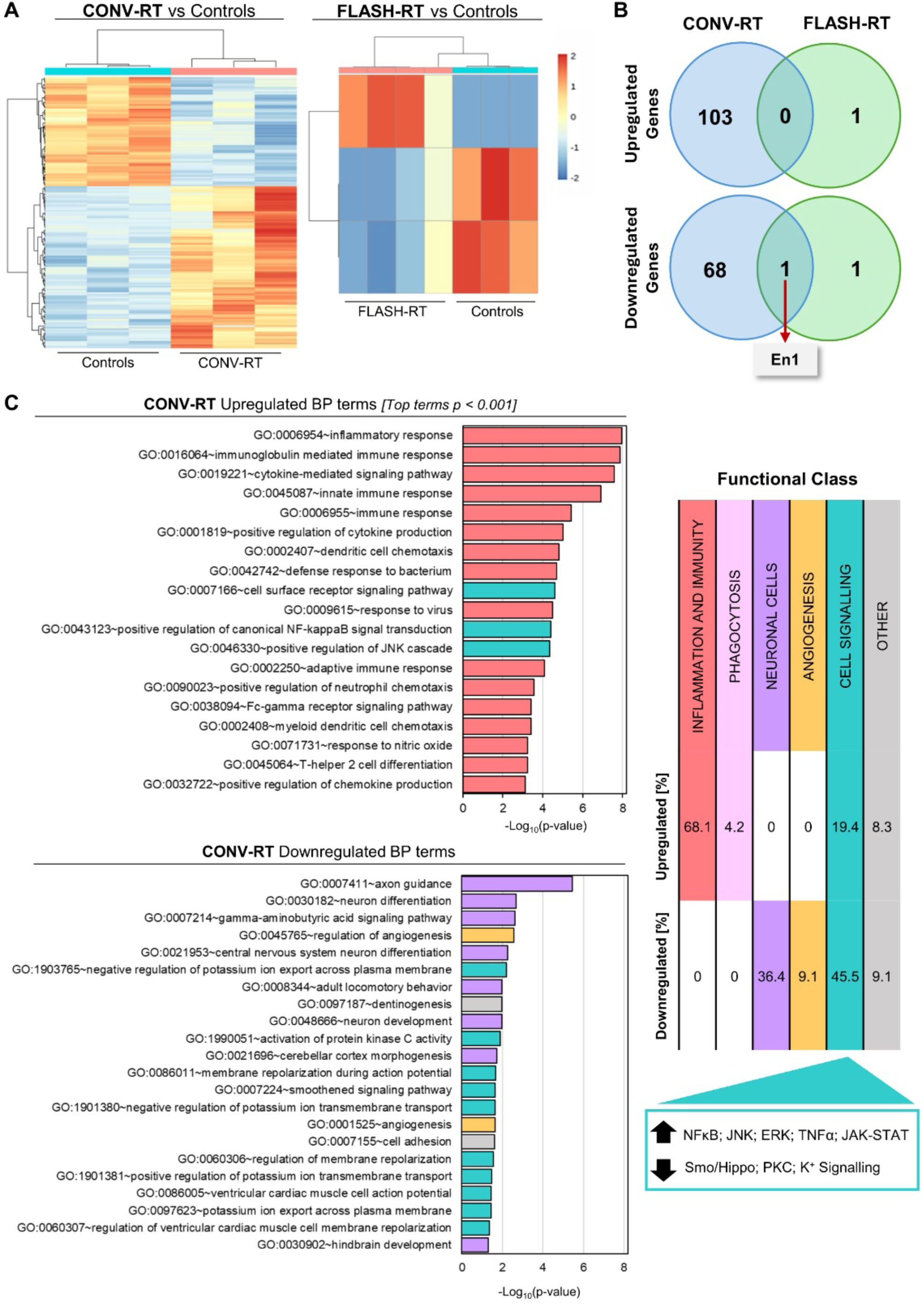
Transcriptomic analysis of left hindlimb iscWAT from mice subjected to CONV- and FLASH-RT. RNA-seq was performed on iscWAT samples collected from the irradiated region of the left hindlimb 70 days after mice received 33.3 Gy electron RT with either CONV-RT or FLASH-RT, as well as from matched non-irradiated CNT (n = 3-4 per group). Differential gene expression analysis among irradiated and CNT groups was conducted using DESeq2. Genes with an absolute fold change (FC) >1.5 and a false discovery rate (FDR)-adjusted p-value<0.05 were considered significantly differentially expressed. **(A)** Heatmaps showing differentially expressed genes in CONV vs. CNT and FLASH vs. CNT comparisons. Color range reflects the log₂ FC between the irradiated group and CNT. **(B)** Venn diagrams representing the number of significantly upregulated and downregulated genes, and the overlap between FLASH-RT and CONV-RT conditions. **(C)** Functional enrichment analysis of GO BP terms among upregulated and downregulated genes in CONV-RT group (p<0.05). Bars represent enriched GO terms ranked by decreasing significance (-log10 p-value) (see also Supplementary Figure 2 for additional significant terms). The various annotation terms collected in the figure are color-coded according to their broad categories, which are detailed in the table. This table also includes the percentages of terms belonging to each category across upregulated and downregulated genes.

By contrast, FLASH-RT, with its minimal effects on iscWAT gene expression, had no clearly enriched biological processes observed.

Collectively, these findings suggest that CONV-RT, but not FLASH-RT, has a profound impact on the gene expression profile of iscWAT. This effect is characterized by the activation of immune and inflammatory processes, including immune cell recruitment, and by the downregulation of pathways essential for proper angiogenesis and neural function.

#### 3.2.3 Histomorphology

To further substantiate the differences between CONV- and FLASH-irradiated iscWAT, we performed additional analyses using light microscopy and TEM. Histology showed signs of altered morphology of adipocytes, suggesting possible suffering and death of these cells. Our and others’ previous work showed that an early sign of suffering and death of adipocytes is the loss of PLIN1, a key protein for the normal functionality of this cell type [41–43]. Quantitative data performed on PLIN1 immunostained sections of iscWAT showed that a large proportion of irradiated adipocytes lost PLIN1 immunoreactivity, in line with the idea that irradiation induces adipocyte suffering and death (Figure 6A). Of note, the effect was more evident in CONV-treated samples (Figure 6A, left panel).

**Figure 6.**
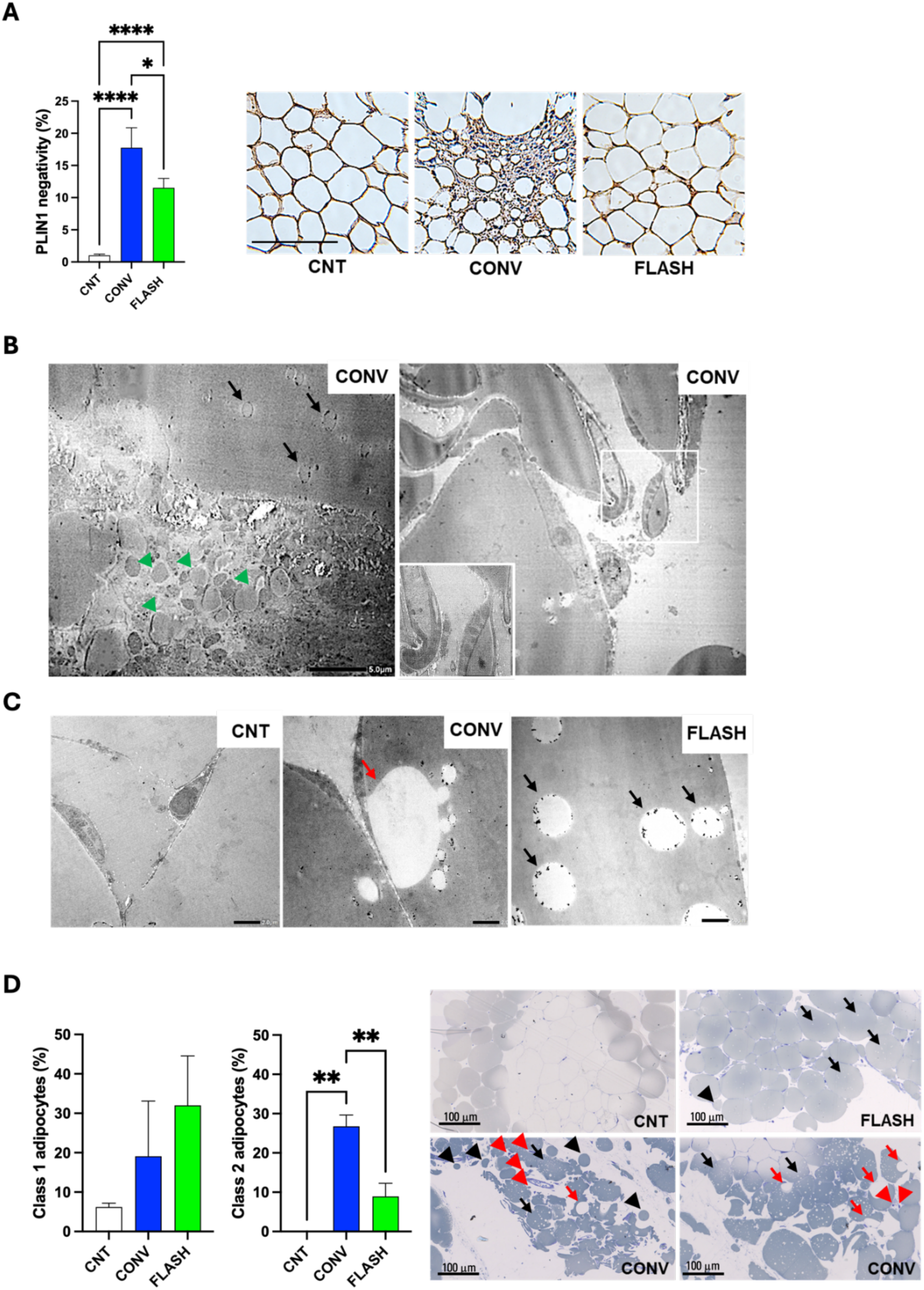
Adipocytes death and signs of damage in irradiated iscWAT. **(A)** Left: Bar graph quantifying the percentage of PLIN1-negative adipocytes. Right: Representative images of iscWAT from CNT, CONV-, and FLASH-irradiated mice immunostained for PLIN1. Scale bar: 100 µm. Magnification: 20x. **(B)** TEM images of iscWAT from CONV-irradiated mice displaying degenerative ultrastructural features. Left: peripheral vacuoles within the adipocyte lipid droplet (black arrows) and extracellular lipid droplets (green arrowheads). Right: irregular, partially delipidated adipocytes (ghost-like structures) with extrusion of droplet-like intracellular material (boxed area, enlarged in inset). Scale bar: 5 µm. Magnification: 1000X. **(C)** TEM images illustrating vacuolization in inguinal scWAT from CNT, CONV-, and FLASH-irradiated mice. CNT adipocytes show homogeneous lipid content with peripheral organelles, whereas irradiated samples display clear vacuoles, predominantly large in CONV (red arrows) and small in FLASH (black arrows). Electron-dense granules at the vacuole periphery represent heavy metal precipitates from the counterstaining procedure. Scale bar: 2 mm. Magnification: 2000X. **(D)** Left: Bar graphs showing morphometric analysis of adipocytes (across the whole section) containing intracellular vacuoles. Adipocytes were classified as Class 1 or Class 2 based on the presence and size of LD-associated vacuoles, with Class 1 presenting at least one vacuole with 2–10 µm major axis and Class 2 presenting at least one vacuole with major axis ≥ 10 µm. Right: Representative toluidine blue–stained semithin sections of iscWAT from CNT, CONV-, and FLASH-irradiated mice. Adipocytes maintain a rounded morphology in CNT and FLASH samples, whereas CONV samples frequently exhibit irregular cell shapes. Extracellular lipid droplets (black arrowheads), small vacuoles (Class 1 adipocytes, black arrows), and large vacuoles (Class 2 adipocytes, red arrows) are indicated. Macrophages are visible in CONV samples (red arrowheads). Scale bar: 100 µm. Magnification: 40X. Data are presented as mean±SEM. Statistical analysis was performed using one-wat ANOVA followed by a multiple-comparison test. * p < 0.05; **p < 0.01; ****p < 0.0001. ANOVA followed by a multiple-comparison test.

In line with these data, TEM showed the presence of structures compatible with degenerating and dead adipocytes (peripheral vacuoles in their lipid droplets, ghost-like adipocytes and free interstitial lipid droplets). Of note, these ultrastructural degenerative aspects were more frequent in CONV treated mice, for which we show representative images in Figures 6B and C. In addition, we noticed the presence of small lipid droplet (LD)-associated vacuoles, that, although present also in control adipocytes, were more prevalent in irradiated iscWAT (Figure 6D, right panel and Suppl. Figure 3). A morphometric analysis of these LD-associated structures was performed by measuring their diameter. Adipocytes were classified into two groups: Class 1 - adipocytes containing at least one vacuole with a major axis of 2–10 μm; Class 2 - adipocytes containing at least one vacuole with a major axis >10 μm. Percentages of Class 1 and 2 adipocytes under the different experimental conditions are shown in the left panel of Figure 6D. Our data indicate a higher number of large LD-associated vacuoles in CONV-RT iscWAT samples.

Death and suffering of adipocytes are accompanied by inflammation mainly due to macrophages, the inflammatory cells responsible for the reabsorption of cellular debris [41] (Figure 7A, right panel, representative TEM image of a macrophage in a CONV-RT sample). In line with an increased damage upon CONV-RT, we found an increased amount of CD68 immunoreactive macrophages in CONV irradiated iscWAT compared to CNT and FLASH-RT samples. (Figure 7A, left panel).

**Figure 7.**
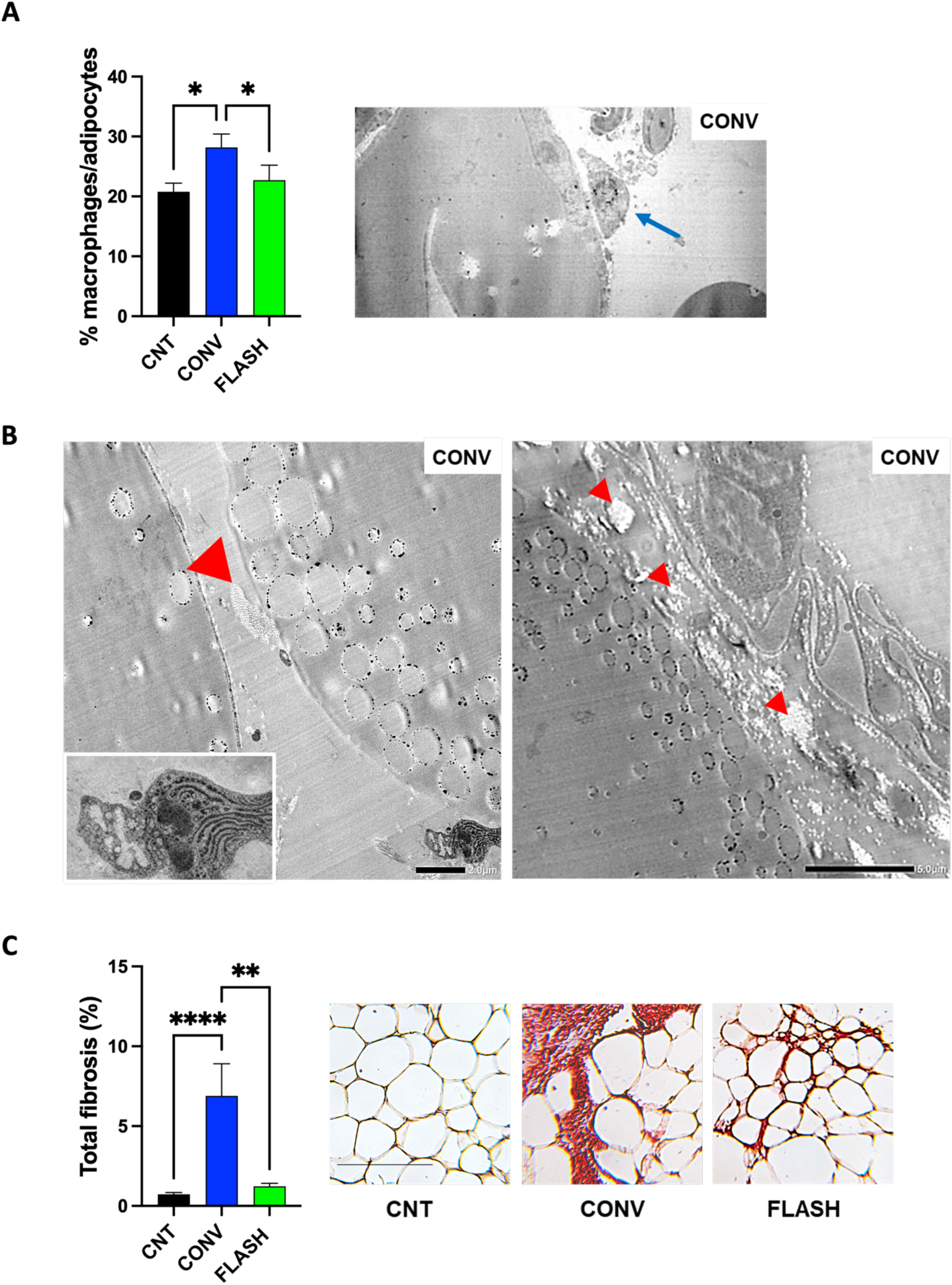
Inflammation and Fibrosis in irradiated iscWAT. **(A)** Left: Bar graph showing the percentage of macrophages relative to total adipocytes in iscWAT from CNT, CONV-, and FLASH-irradiated mice (≥ 40 fields analyzed per group). Right: Representative TEM image of CONV-irradiated inguinal scWAT; the blue arrow indicates a macrophage in close proximity to an adipocyte. Scale bar: 5 µm. Magnification: 1000X. **(B)** Representative TEM images of iscWAT from CONV-irradiated mice showing fibrotic features, evidenced by extracellular collagen accumulation (red arrowheads). An activated fibroblast with abundant rough endoplasmic reticulum (RER) is visible in the left image and enlarged in the inset. Scale bars: 2 µm (left) and 5 µm (right). Magnification: 2000× (left) and 1500× (right). **(C)** Left: Bar graph quantifying the percentage of fibrotic area in iscWAT from CNT, CONV-, and FLASH-irradiated mice. Right: Representative Sirius Red–stained sections of inguinal scWAT from CNT, CONV-, and FLASH-irradiated mice highlighting collagen deposition. Scale bar: 100 µm. Magnification: 20x. Data are presented as mean±SEM. Statistical analysis was performed using Kruskal–Wallis test followed by multiple-comparison tests. *p < 0.05; **p < 0.01; ****p < 0.0001.

Fibrosis is a common finding of irradiated tissues [48,49]. In line, we found ultrastructural evidence of activated fibroblasts in irradiated iscWAT (Figure 7B). Sirius red staining is the gold standard method for histological quantification of fibrosis [40]; our quantitative analysis of Sirius Red–stained sections showed that fibrosis was present in irradiated adipose tissue, with higher levels observed in CONV-treated iscWAT (Figure 7C). According to our data, the skin of irradiated mice is characterized by increased thickness, a hallmark of fibrosis [19]. We next sought to determine whether the signs of distress observed in the subcutaneous WAT, the hypodermal layer, were associated with changes in skin thickness. Indeed, we found a highly significant and positive correlation between the proportion of PLIN1-negative cells and both epidermal and dermal thickness (Suppl. Figure 4), indicating that these represent two distinct but related aspects of tissue response to RT.

Collectively, these findings support the notion that CONV-RT induces more severe damage in healthy scWAT than FLASH-RT.

## 4. Discussion

Subcutaneous white adipose tissue (WAT) accounts for approximately 30–60% of total body weight, depending on the degree of obesity [50]. Beyond serving as the principal energy reservoir in mammals, WAT functions as a major endocrine organ regulating appetite, insulin sensitivity, systemic inflammation [44], and bone turnover [51]. In addition, WAT contributes to immune modulation and provides mechanical protection. Importantly, adipocytes are frequently located in close proximity to cancer cells in many solid tumors [52].

Given these functions, WAT represents an unavoidable target in cancer patients undergoing RT, and radiation-induced alterations in its structure or function may have consequences extending far beyond the irradiated area. Nevertheless, current RT planning strategies do not consider potential collateral damage to WAT. A PubMed search combining the terms “adipose tissue” and “radiotherapy” retrieves only a few hundred publications, most (>80%) focusing on the relationship between obesity and cancer prognosis or on the use of adipose-derived stem cells for tissue repair after cancer or RT-induced damage. When the search is refined to “FLASH radiotherapy” and “adipose tissue”, no results are retrieved, underscoring the absence of data in this emerging field. In this context, it is striking that the consequences of ionizing radiation on WAT function remain poorly investigated [6,7]. Indirect evidence nevertheless suggests that WAT is sensitive to radiation exposure. Cross-sectional studies of individuals who received RT during childhood show significant differences in metabolic parameters compared with untreated individuals, and RT has also been implicated in cases of acquired localized lipodystrophy. Together, these observations support the notion that WAT responds to ionizing radiation with consequences that may extend beyond the irradiated field.

In the present study, we outline a stepwise trajectory describing how RT may affect WAT, progressing from cellular alterations to organ-level dysfunction. Importantly, we show that when RT is delivered at UHDR, this trajectory is interrupted or markedly attenuated.

The first prerequisite for WAT maintenance is the preservation of a healthy pool of preadipocytes within the basal lamina niche, which provides a reservoir of proliferative progenitors [53]. Approximately 10% of adipocytes are renewed annually in humans through differentiation of these progenitors [54]. Our data indicate that neither CONV- nor FLASH-RT significantly affect preadipocyte viability, suggesting that the progenitor reservoir remains largely preserved, at least in the short term.

The second step in this trajectory is adipogenesis. Physiological renewal of WAT relies on differentiation of preadipocytes into mature adipocytes to replace cells lost over time. *In vitro*, RT markedly impairs adipogenesis in a dose-dependent manner, consistent with previous reports [55]. At this stage, differences between RT modalities become evident: FLASH-RT preserves adipogenic capacity more effectively than CONV-RT at low-to-intermediate doses, suggesting partial protection of tissue regenerative potential under UHDR conditions.

The third step concerns the response of mature adipocytes. Although RT has limited effects on adipocyte viability —and negligible effects after FLASH-RT— it induces a pronounced senescence phenotype. Again, UHDR exposure results in a comparatively attenuated response, whereas CONV-RT triggers marked senescence. Accumulation of senescent adipocytes is expected to promote chronic tissue dysfunction through altered secretory profiles and impaired metabolic regulation, potentially leading to systemic consequences. Consistent with this interpretation, we previously showed that CONV-, but not FLASH-RT, alters appetite-regulating hormones and increases circulating inflammatory markers [19].

The fourth step concerns the *in vivo* impact of RT on WAT. Interestingly, we observed not only a reduction in irradiated fat pads but also in WAT depots that were not directly targeted by irradiation. This finding suggests systemic effects and/or inter-depot communication mechanisms, potentially mediated by extracellular vesicles and circulating nucleic acids [56]. Such mechanisms may explain why WAT, although anatomically dispersed throughout the body, functions as a coordinated endocrine organ capable of mounting integrated responses to external insults [57]. Within this framework, local radiation damage may trigger systemic signaling affecting distant fat depots. Consistent with this model, transcriptomic and functional enrichment analyses indicate that CONV-RT, but not FLASH-RT, induces a robust inflammatory response in iscWAT while suppressing biological processes associated with normal adipose function. The pronounced transcriptional alterations observed after CONV-RT, compared with the negligible changes following FLASH-RT, closely mirror our previous observations in murine skin and skeletal muscle [19], suggesting that this differential tissue response may represent a broader biological feature of CONV versus FLASH irradiation. Molecular alterations are paralleled by persistent structural changes. IscWAT exposed to CONV-RT displays markedly more severe morphological abnormalities than FLASH-treated tissue, including vacuolization of lipid droplets, extrusion of lipid material into the extracellular space, and the presence of “ghost” adipocytes. These features are consistent with progressive adipocyte degeneration and suggest engagement of a potentially irreversible pathway leading to cell death. Supporting this interpretation, the loss of perilipin immunoreactivity observed in CONV-irradiated tissue indicates extensive adipocyte loss or suffering. Notably, these findings in hypodermal WAT are consistent with the markedly reduced expression of key adipose markers—perilipin, leptin, and adiponectin—that we previously described in intradermal WAT [19]. Together, these observations support the concept that RT disrupts adipose tissue homeostasis at both structural and molecular levels.

Our results align with previous reports describing WAT dysfunction in adults who received RT during childhood [18,58] and suggest a trajectory through which the long-term consequences of RT may culminate in lipodystrophy. Indeed, localized lipodystrophy has been reported years after RT exposure [59], and there is a striking similarity between certain laminopathic forms of lipodystrophy and those acquired after RT [60], particularly in terms of their characteristic fat redistribution.

We propose that this process develops slowly and progressively: adipocytes that die are not adequately replaced by newly differentiated cells, while chronic inflammation and fibrosis remodel the tissue. Over time, adipose tissue may be replaced by a non-specialized connective matrix, giving rise to tissue unable to store triglycerides and to properly sense and dispose of glucose.

## Conclusions

This study identifies white adipose tissue as a previously overlooked target of ionizing radiation and outlines a trajectory through which RT may progressively impair WAT homeostasis. From altered adipogenesis and induction of adipocyte senescence to systemic inflammatory responses and structural degeneration of fat depots, our findings suggest that radiation exposure can initiate a cascade of events that ultimately compromises WAT function and lead to lipodystrophy. Importantly, when RT is delivered at ultra-high dose rate, many of these alterations are markedly attenuated. FLASH-RT preserves adipogenic potential, limits senescence, reduces inflammatory transcriptional responses, and largely prevents the structural degeneration observed after conventional irradiation. Together, these findings indicate that adipose tissue may represent an additional normal-tissue compartment benefiting from the FLASH effect. Given the central metabolic and endocrine functions of WAT, its preservation may have implications extending beyond the irradiated field, potentially influencing systemic metabolism and long-term health in cancer survivors.

## Limitations

First, the study was conducted primarily in murine models and *in vitro* systems, and extrapolation to human WAT physiology should therefore be made with caution. Second, although our experiments capture early and intermediate responses to irradiation, longer follow-up will be necessary to determine whether the observed alterations ultimately translate into clinically relevant metabolic dysfunction or lipodystrophic phenotypes. Third, the complete molecular mechanisms underlying the differential response of WAT to CONV and FLASH irradiation remain to be elucidated, as while we identify genes and pathways consistent with the for the observed protective effect under UHDR conditions, the upstream common orchestrator sitting at the interaction between the physical impulse (electrons) and the start of the biological cascade remain to be elucidated.

Future studies combining long-term metabolic analyses, mechanistic investigations, and clinical observations will be necessary to clarify the role of WAT in radiation-induced normal tissue toxicity and to determine whether the protective effects of FLASH-RT observed here extend to human patients.

## Acknowledgments

We acknowledge Fondazione Pisa for funding CPFR with the grant “prog. n.134/2021” and the Federal Ministry of Research, Technology and Space (Bundesministerium für Forschung, Technologie und Raumfahrt, BMFTR) as part of the German Center for Child and Adolescent Health (DZKJ) for funding Martin Wabitsch (funding code 01GL2407A).

A special thanks to Emma Buzzigoli for technical assistance and to Cecilia Ciampi, Sara Ciampi and Gianpietro Chessa for the daily management of animals.

## Funding

· Piano Nazionale di Ripresa e Resilienza (PNRR), Missione 4, Componente 2, Ecosistemi dell’Innovazione–Tuscany Health Ecosystem (THE), Spoke 1 “Advanced Radiotherapies and Diagnostics in Oncology”—CUP I53C22000780001 (MM) and cascading call financed grant RADAR.

· PNRR-MAD-2022-12376459, CUP B53C22009530001 (MM)

· MIRO - MInibeam RadiOtherapy Program: INFN CSN5 Call 2024-2026 (MM)

· AIRC - IG 2024 ID-30434 project (MC)

## Author contributions: CrediT

**Conceptualization:** MM, SCi, FDM

**Data curation:** GS, GF, EMDS, MM

**Formal analysis:** GS, GF, CK, AC, FR, RDP, EG, SL, TL, RC, SCi, FDM, MM

**Funding acquisition:** MM, SCas, SCap, FDM, FP

**Investigation:** GS, GF, EMDS, FR, SL, TL, MM

**Methodology:** GS, GF, AU, GA, EG, EMDS, CK, AC, FR, RDP, SL, TL, RC, SCi, AG, FP, FDM, MM

**Project administration:** MM, SCas

**Resources:** MM, SCas, SCap, FP, FDM, MW, MC, SCi

**Supervision:** MM, FDM, SCi, CK

**Validation:** GS, SCi, MM

**Visualization:** GS, GF, EMDS, CK, RDP, SL, TL, SCi, MM

**Writing – original draft**: MM

**Writing – review & editing:** MM, GS, GF, AU, GA, CK, AC, RDP, SL, TL, RC, SCi, FDM, AG, FP

## Declaration of generative AI and AI-assisted technologies in the manuscript preparation process

During the preparation of this work the authors used ChatGPT in order to assist with grammar and language. After using this tool, the authors reviewed and edited the content as needed and take full responsibility for the content of the published article.

## Abbreviations

ADR: Average Dose Rate
BP: Biological Process
CNT: Control
CONV-RT: Conventional radiotherapy
DPP: Dose Per Pulse
EPI WAT: Epididymal White Adipose Tissue
FLASH-RT: Ultra-high dose rate (FLASH) radiotherapy
GO: Gene Ontology
IR: Ionizing Radiation
iscWAT: Inguinal Subcutaneous White Adipose Tissue
LD: Lipid Droplets
LINAC: Linear Accelerator
PERI WAT: Perirenal White Adipose Tissue
PID: Post-Induction Day
PRF: Pulse Repetition Frequency
RT: Radiotherapy
scWAT: Subcutaneous White Adipose Tissue
SGBS: Simpson–Golabi–Behmel Syndrome (cell line)
TG: Triglycerides
WAT: White Adipose Tissue

