## Supplementary Figures and Tables for "Differential impact of FLASH and conventional radiotherapy on a pivotal metabolic organ: White Adipose Tissue"

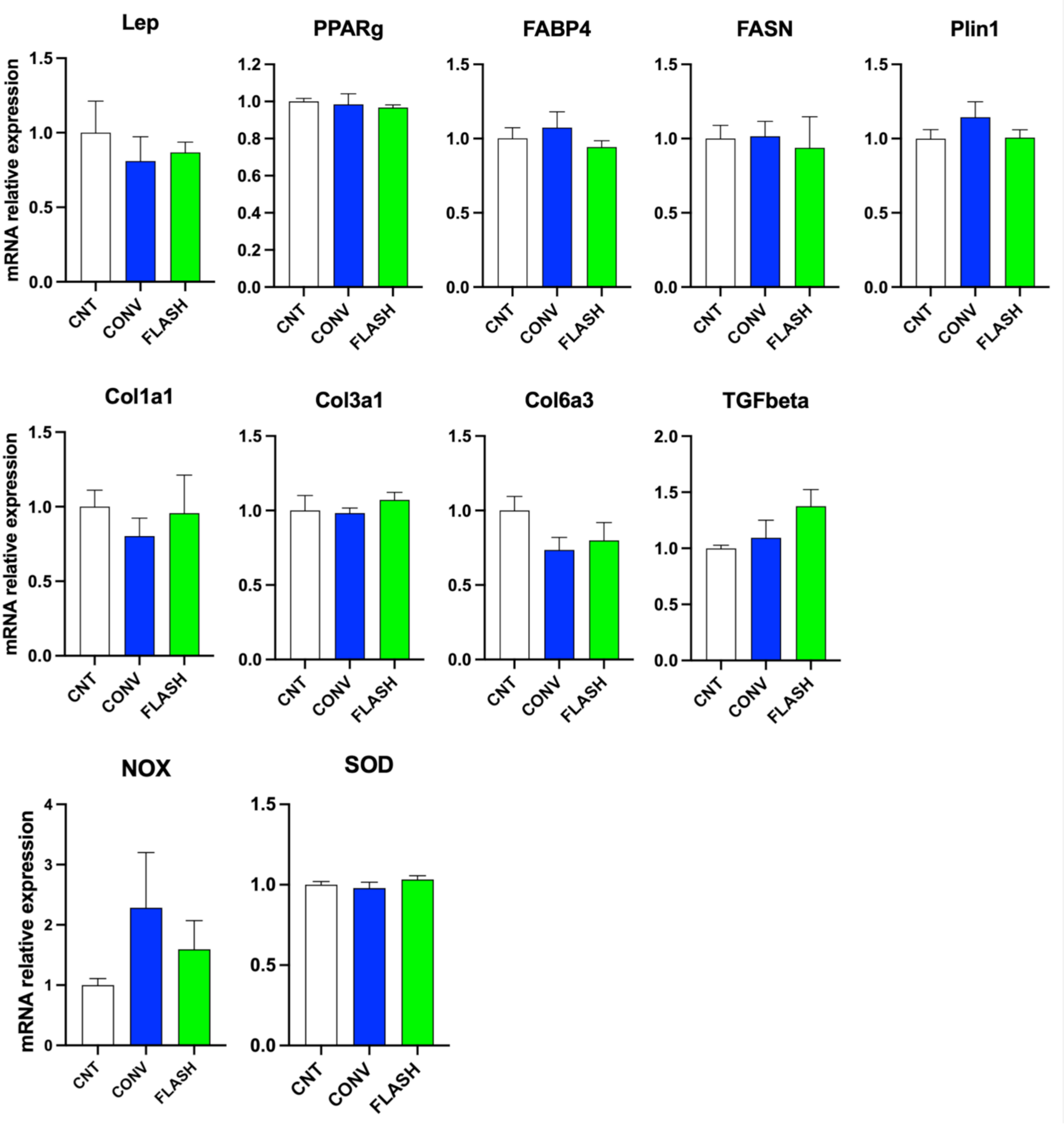


**Supplementary Figure 1. Gene expression analysis on CONV- and FLASH-irradiated adipocytes: adipocyte maturation, fibrosis and oxidative stress.** Gene expression analysis of genes associated with adipocyte maturation (LEP, PPARG, FABP4, FASN and PLIN1), fibrosis (Col1a1, Col3a1, Col6a3 and TGFbeta), and oxidative stress generation and detoxification (NOX and SOD). Data are presented as mean ± SEM. Statistical analysis was performed using one-way ANOVA or Kruskal–Wallis test, as appropriate.

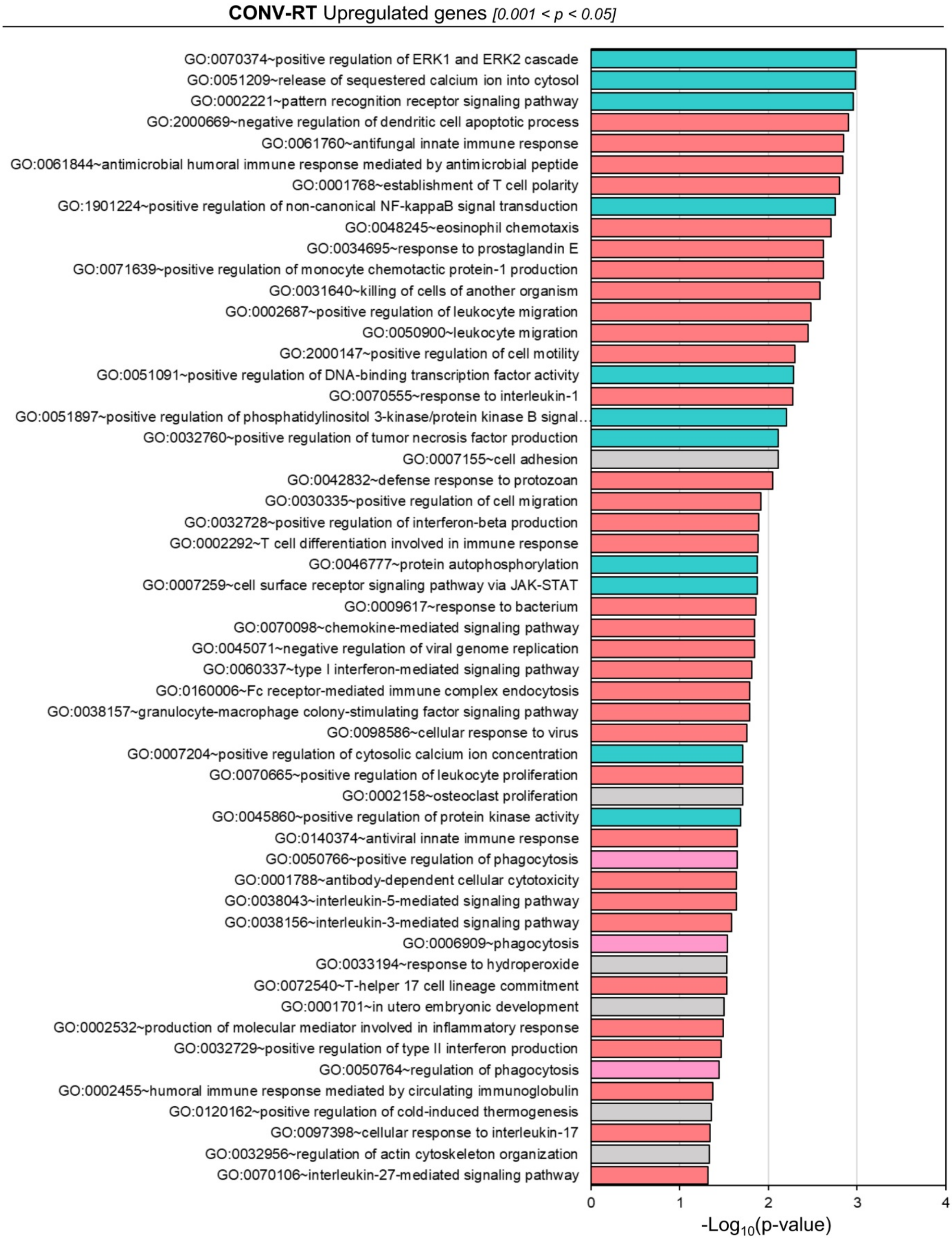


**Supplementary Figure 2. Transcriptomic analysis of left hindlimb iscWAT tissue from mice subjected to CONV- and FLASH-RT, supplementary analysis.** RNA-seq was performed on iscWAT samples collected from the irradiated region of the left hindlimb 70 days after mice received 33.3 Gy electron RT with either CONV-RT or FLASH-RT, as well as from matched non-irradiated CNT (n = 3-4 per group). Differential gene expression analysis among irradiated and CNT groups was conducted using DESeq2. Genes with an absolute fold change >1.5 and a false discovery rate (FDR)-adjusted p-value<0.05 were considered significantly differentially expressed. Functional enrichment analysis of Gene Ontology (GO) biological process (BP) terms among upregulated and downregulated genes in CONV-RT group (p<0.05). Bars represent enriched GO BPterms ranked by decreasing significance (-log10 p-value). The main data are shown in Figure 5C; this figure represents additional enriched terms among the CONV-RT upregulated genes, 0.001 < p-value < 0.05.  The various annotation terms collected in the figure are color-coded according to their broad categories, which are detailed in the table presented in Figure 5C.

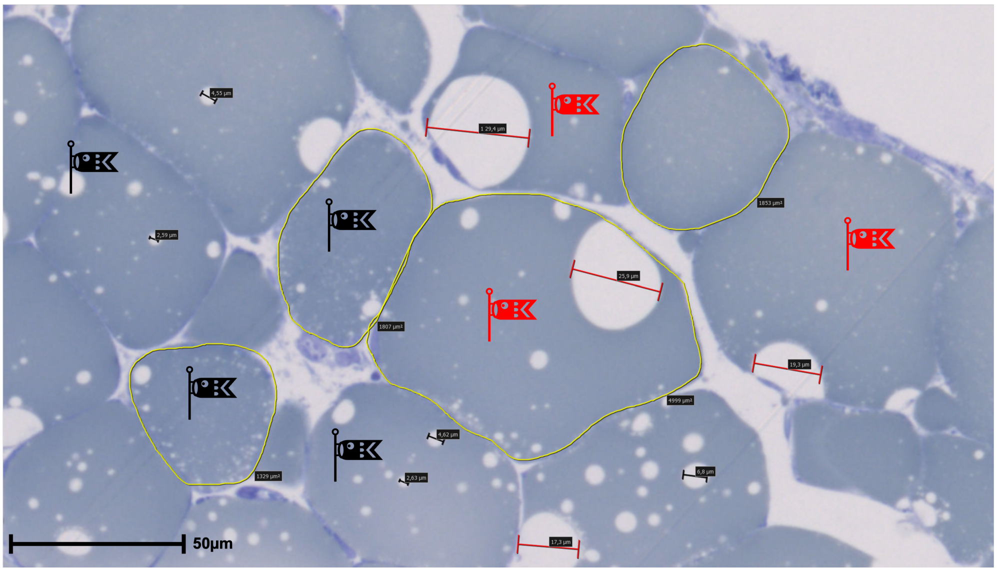


**Supplementary Figure 3. Morphometric analysis on toluidine blue-stained semithin sections of CONV-irradiated murine inguinal subcutaneous WAT**. Morphometry was performed on all sections with the Hamamatsu NDP view 2.0 software (see Methods). The picture shows examples of evaluation of adipocyte area (yellow profiles) and vacuole size (black bar: small vacuoles, major axis 2–10 µm; red bar: large vacuoles, major axis >10 µm) with actual measures indicated in the labels. Class 1 adipocytes (presenting at least one small vacuole) are marked with a black flag, Class 2 adipocytes (presenting at least one large vacuole) are marked with a red flag. Scale bar: 50 µm. Magnification: 80X.

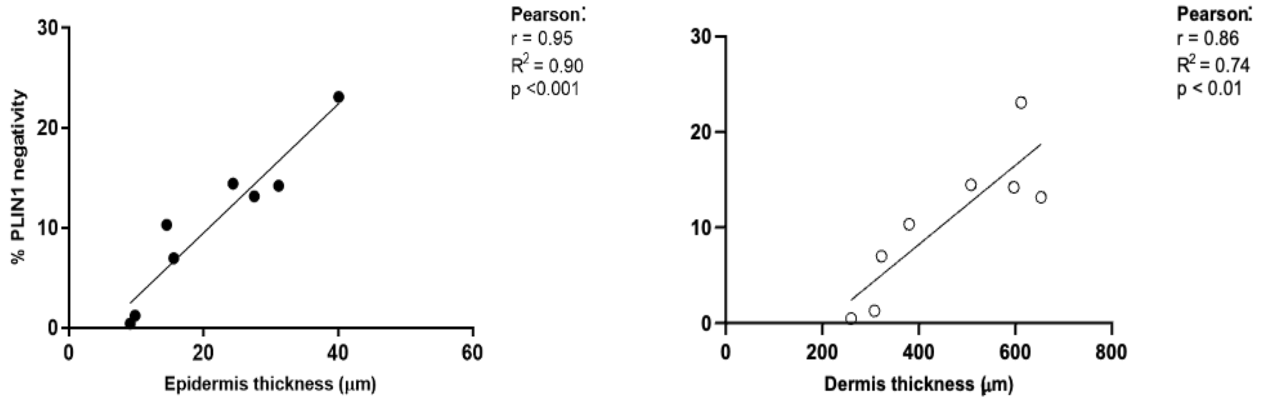


**Supplementary Figure 4.** Correlation plots illustrating the relationship between epidermis or demis thickness (as indicated) and PLIN negativity in the intradermal adipose tissue underneath.

### Supplementary Tables

**Supplementary Table 1: Upregulated genes in CONV-RT mice iscWAT compared to CNT.** Genes with fold change >1.5 and a false discovery rate (FDR)-adjusted p-value<0.05 were considered significantly upregulated.

| Gene Name | Description | log2(FC) | pvalue | padj |
|---|---|---|---|---|
| **Socs3** | suppressor of cytokine signaling 3 | **2.70** | 3.79E-23 | **3.70E-19** |
| **Ccl8** | C-C motif chemokine ligand 8 | **3.54** | 1.63E-11 | **5.31E-08** |
| **Sirpb1a** | signal-regulatory protein beta 1A | **3.64** | 2.79E-11 | **7.79E-08** |
| **Lcn2** | lipocalin 2 | **4.98** | 1.39E-10 | **3.38E-07** |
| **Sirpb1c** | signal-regulatory protein beta 1C | **3.04** | 7.21E-10 | **1.41E-06** |
| **Mcoln2** | mucolipin 2 | **3.17** | 1.50E-09 | **2.66E-06** |
| **Pgap6** | post-glycosylphosphatidylinositol attachment to proteins 6 | **1.27** | 4.43E-09 | **7.21E-06** |
| **Hs3st3b1** | heparan sulfate (glucosamine) 3-O-sulfotransferase 3B1 | **2.06** | 2.41E-08 | **2.94E-05** |
| **Il4ra** | interleukin 4 receptor, alpha | **1.82** | 3.65E-08 | **3.96E-05** |
| **Iglc2** | immunoglobulin lambda constant 2 | **7.33** | 5.99E-08 | **6.15E-05** |
| **Chil3** | chitinase-like 3 | **5.87** | 2.03E-07 | **1.72E-04** |
| **Clec4n** | C-type lectin domain family 4, member n | **2.47** | 2.85E-07 | **2.22E-04** |
| **Ighv9-3** | immunoglobulin heavy variable V9-3 | **13.67** | 3.46E-07 | **2.50E-04** |
| **Oas2** | 2'-5' oligoadenylate synthetase 2 | **1.84** | 4.69E-07 | **3.23E-04** |
| **Zfp990** | zinc finger protein 990 | **5.21** | 1.03E-06 | **5.93E-04** |
| **Serpina3i** | serine (or cysteine) peptidase inhibitor, clade A, member 3I | **3.41** | 1.02E-06 | **5.93E-04** |
| **Rbm20** | RNA binding motif protein 20 | **2.03** | 1.34E-06 | **7.08E-04** |
| **Pirb** | paired Ig-like receptor B | **1.28** | 1.41E-06 | **7.25E-04** |
| **Ripor3** | RIPOR family member 3 | **2.60** | 1.46E-06 | **7.31E-04** |
| **Fcgr2b** | Fc receptor, IgG, low affinity IIb | **1.60** | 1.55E-06 | **7.57E-04** |
| **Bcl3** | B cell leukemia/lymphoma 3 | **2.16** | 2.27E-06 | **1.05E-03** |
| **Mefv** | Mediterranean fever | **3.53** | 2.66E-06 | **1.21E-03** |
| **Ccl19-ps6** | C-C motif chemokine ligand 19, pseudogene 6 | **3.66** | 2.91E-06 | **1.29E-03** |
| **Igkc** | immunoglobulin kappa constant | **8.60** | 3.55E-06 | **1.45E-03** |
| **Gm5150** | predicted gene 5150 | **3.14** | 5.01E-06 | **2.00E-03** |
| **Fbn2** | fibrillin 2 | **1.74** | 5.46E-06 | **2.09E-03** |
| **Nectin4** | nectin cell adhesion molecule 4 | **1.65** | 6.33E-06 | **2.38E-03** |
| **Igha** | immunoglobulin heavy constant alpha | **4.09** | 6.91E-06 | **2.54E-03** |
| **Ccl19-ps5** | C-C motif chemokine ligand 19, pseudogene 5 | **6.72** | 7.18E-06 | **2.59E-03** |
| **Igkv15-103** | immunoglobulin kappa chain variable 15-103 | **10.09** | 7.47E-06 | **2.65E-03** |
| **Plac8** | placenta-specific 8 | **2.64** | 7.72E-06 | **2.69E-03** |
| **Il1b** | interleukin 1 beta | **3.17** | 8.19E-06 | **2.81E-03** |
| **Oas1g** | 2'-5' oligoadenylate synthetase 1G | **1.51** | 9.83E-06 | **3.25E-03** |
| **Lipg** | lipase, endothelial | **3.02** | 1.13E-05 | **3.68E-03** |
| **Dnah6** | dynein, axonemal, heavy chain 6 | **1.97** | 1.20E-05 | **3.82E-03** |
| **Slfn1** | schlafen 1 | **3.37** | 1.82E-05 | **5.23E-03** |
| **Sirpd** | signal regulatory protein delta | **3.91** | 1.98E-05 | **5.61E-03** |
| **Slfn4** | schlafen 4 | **3.30** | 2.02E-05 | **5.61E-03** |
| **Ccl12** | C-C motif chemokine ligand 12 | **2.47** | 2.04E-05 | **5.61E-03** |
| **Cd38** | CD38 antigen | **1.65** | 2.20E-05 | **5.88E-03** |
| **Sspo** | SCO-spondin | **1.61** | 2.51E-05 | **6.54E-03** |
| **Clec4d** | C-type lectin domain family 4, member d | **4.39** | 2.56E-05 | **6.58E-03** |
| **Csf3r** | colony stimulating factor 3 receptor | **2.40** | 2.94E-05 | **7.27E-03** |
| **Gpr141** | G protein-coupled receptor 141 | **2.71** | 3.47E-05 | **7.89E-03** |
| **Ccl19** | C-C motif chemokine ligand 19 | **2.25** | 3.39E-05 | **7.89E-03** |
| **Gm18988** | predicted gene, 18988 | **1.89** | 3.31E-05 | **7.89E-03** |
| **Igkv8-30** | immunoglobulin kappa chain variable 8-30 | **7.45** | 3.57E-05 | **8.02E-03** |
| **Fpr1** | formyl peptide receptor 1 | **5.34** | 4.04E-05 | **8.85E-03** |
| **Fpr2** | formyl peptide receptor 2 | **4.11** | 5.44E-05 | **1.15E-02** |
| **Hck** | hemopoietic cell kinase | **2.80** | 5.48E-05 | **1.15E-02** |
| **Spon1** | spondin 1, (f-spondin) extracellular matrix protein | **1.06** | 5.78E-05 | **1.20E-02** |
| **Fcgr4** | Fc receptor, IgG, low affinity IV | **2.27** | 6.46E-05 | **1.30E-02** |
| **Zdhhc23** | zinc finger, DHHC domain containing 23 | **1.62** | 6.37E-05 | **1.30E-02** |
| **Fgr** | FGR proto-oncogene, Src family tyrosine kinase | **3.13** | 6.79E-05 | **1.35E-02** |
| **Pla2g7** | phospholipase A2, group VII (platelet-activating factor acetylhydrolase, plasma) | **1.71** | 7.00E-05 | **1.38E-02** |
| **Sele** | selectin, endothelial cell | **2.18** | 7.41E-05 | **1.45E-02** |
| **Trpm2** | transient receptor potential cation channel, subfamily M, member 2 | **3.26** | 7.89E-05 | **1.52E-02** |
| **Emilin1** | elastin microfibril interfacer 1 | **0.96** | 8.15E-05 | **1.55E-02** |
| **Ceacam16** | CEA cell adhesion molecule 16 | **5.13** | 8.63E-05 | **1.57E-02** |

| | | | | |
|---|---|---|---|---|
| **Chl1** | cell adhesion molecule L1-like | **2.58** | 8.54E-05 | **1.57E-02** |
| **Pim1** | proviral integration site 1 | **1.90** | 8.54E-05 | **1.57E-02** |
| **Clec4e** | C-type lectin domain family 4, member e | **2.17** | 8.97E-05 | **1.62E-02** |
| **Junb** | jun B proto-oncogene | **1.28** | 9.84E-05 | **1.75E-02** |
| **Ighv14-2** | immunoglobulin heavy variable 14-2 | **9.84** | 1.01E-04 | **1.78E-02** |
| **Itih3** | inter-alpha trypsin inhibitor, heavy chain 3 | **3.03** | 1.10E-04 | **1.88E-02** |
| **B4galt5** | UDP-Gal:betaGlcNAc beta 1,4-galactosyltransferase, polypeptide 5 | **1.45** | 1.09E-04 | **1.88E-02** |
| **Ms4a4a** | membrane-spanning 4-domains, subfamily A, member 4A | **1.74** | 1.13E-04 | **1.92E-02** |
| **Tnfsf11** | tumor necrosis factor (ligand) superfamily, member 11 | **5.13** | 1.22E-04 | **2.04E-02** |
| **Tlr13** | toll-like receptor 13 | **1.73** | 1.43E-04 | **2.37E-02** |
| **Mid1** | midline 1 | **1.06** | 1.50E-04 | **2.46E-02** |
| **Igkv2-112** | immunoglobulin kappa variable 2-112 | **7.64** | 1.52E-04 | **2.48E-02** |
| **Siglec1** | sialic acid binding Ig-like lectin 1, sialoadhesin | **1.19** | 1.55E-04 | **2.50E-02** |
| **Entpd3** | ectonucleoside triphosphate diphosphohydrolase 3 | **3.08** | 1.58E-04 | **2.52E-02** |
| **Batf** | basic leucine zipper transcription factor, ATF-like | **2.66** | 1.63E-04 | **2.54E-02** |
| **Gm867** | predicted gene 867 | **3.90** | 1.76E-04 | **2.64E-02** |
| **Ms4a6c** | membrane-spanning 4-domains, subfamily A, member 6C | **2.12** | 1.74E-04 | **2.64E-02** |
| **Csf2rb2** | colony stimulating factor 2 receptor, beta 2, low-affinity (granulocyte-macrophage) | **1.75** | 1.75E-04 | **2.64E-02** |
| **Runx1** | runt related transcription factor 1 | **1.58** | 1.77E-04 | **2.64E-02** |
| **Ifitm1** | interferon induced transmembrane protein 1 | **2.95** | 1.83E-04 | **2.66E-02** |
| **Ly6a** | lymphocyte antigen 6 family member A | **0.79** | 1.84E-04 | **2.66E-02** |
| **Ms4a6b** | membrane-spanning 4-domains, subfamily A, member 6B | **2.60** | 1.95E-04 | **2.74E-02** |
| **Csf2rb** | colony stimulating factor 2 receptor, beta, low-affinity (granulocyte-macrophage) | **1.20** | 1.94E-04 | **2.74E-02** |
| **Sik1** | salt inducible kinase 1 | **0.90** | 2.04E-04 | **2.84E-02** |
| **Ighv5-17** | immunoglobulin heavy variable 5-17 | **10.23** | 2.31E-04 | **3.20E-02** |
| **Tent5c** | terminal nucleotidyltransferase 5C | **2.54** | 2.35E-04 | **3.23E-02** |
| **Susd1** | sushi domain containing 1 | **1.35** | 2.60E-04 | **3.50E-02** |
| **Madcam1** | mucosal vascular addressin cell adhesion molecule 1 | **5.85** | 2.76E-04 | **3.66E-02** |
| **Cd300c2** | CD300C molecule 2 | **1.53** | 2.87E-04 | **3.76E-02** |
| **Flvcr1** | feline leukemia virus subgroup C cellular receptor 1 | **0.73** | 2.97E-04 | **3.84E-02** |
| **Lrrc25** | leucine rich repeat containing 25 | **1.07** | 3.17E-04 | **4.06E-02** |
| **Amd1** | S-adenosylmethionine decarboxylase 1 | **0.64** | 3.18E-04 | **4.06E-02** |
| **Ighv1-72** | immunoglobulin heavy variable 1-72 | **8.44** | 3.31E-04 | **4.17E-02** |
| **Ly9** | lymphocyte antigen 9 | **3.25** | 3.29E-04 | **4.17E-02** |
| **H2bc24** | H2B clustered histone 24 | **3.06** | 3.37E-04 | **4.18E-02** |
| **Cyp7b1** | cytochrome P450, family 7, subfamily b, polypeptide 1 | **1.83** | 3.36E-04 | **4.18E-02** |
| **Igkv8-28** | immunoglobulin kappa variable 8-28 | **9.80** | 3.51E-04 | **4.30E-02** |
| **Pira1** | paired-Ig-like receptor A1 | **2.18** | 4.00E-04 | **4.74E-02** |
| **Man2a1** | mannosidase 2, alpha 1 | **0.63** | 4.03E-04 | **4.74E-02** |
| **Krt5** | keratin 5 | **5.12** | 4.07E-04 | **4.75E-02** |
| **Cpxm1** | carboxypeptidase X, M14 family member 1 | **1.31** | 4.16E-04 | **4.84E-02** |
| **Slc25a37** | solute carrier family 25, member 37 | **1.17** | 4.28E-04 | **4.90E-02** |
| **Rgs2** | regulator of G-protein signaling 2 | **1.16** | 4.30E-04 | **4.90E-02** |
| **Sting1** | stimulator of interferon response cGAMP interactor 1 | **2.01** | 4.33E-04 | **4.91E-02** |

**Supplementary Table 2: Downregulated genes in CONV-RT mice iscWAT compared to CNT.** Genes with fold change < -1.5 and a false discovery rate (FDR)-adjusted p-value<0.05 were considered significantly downregulated.

| Gene Name | Description | log2(FC) | pvalue | padj |
|---|---|---|---|---|
| **Prnd** | prion like protein doppel | **-2.75** | 9.30E-31 | **1.81E-26** |
| **D430019H16Rik** | RIKEN cDNA D430019H16 gene | **-3.02** | 1.53E-20 | **9.93E-17** |
| **Gpr156** | G protein-coupled receptor 156 | **-2.10** | 6.03E-19 | **2.94E-15** |
| **Rgs7bp** | regulator of G-protein signalling 7 binding protein | **-2.47** | 1.10E-13 | **4.31E-10** |
| **Ankrd29** | ankyrin repeat domain 29 | **-2.62** | 5.23E-10 | **1.13E-06** |
| **Mest** | mesoderm specific transcript | **-3.72** | 1.06E-08 | **1.59E-05** |
| **Kcne5** | potassium voltage-gated channel subfamily E regulatory subunit 5 | **-3.05** | 1.76E-08 | **2.40E-05** |
| **Bpifb6** | BPI fold containing family B, member 6 | **-3.70** | 1.84E-08 | **2.40E-05** |
| **Hpd** | 4-hydroxyphenylpyruvic acid dioxygenase | **-2.26** | 2.59E-08 | **2.97E-05** |
| **Unc5b** | unc-5 netrin receptor B | **-1.08** | 6.91E-08 | **6.75E-05** |
| **Akr1c14** | aldo-keto reductase family 1, member C14 | **-1.45** | 1.14E-07 | **1.01E-04** |
| **Prss51** | serine protease 51 | **-4.63** | 1.09E-07 | **1.01E-04** |
| **Serpina1c** | serine (or cysteine) peptidase inhibitor, clade A, member 1C | **-2.67** | 2.44E-07 | **1.98E-04** |
| **Nkd1** | naked cuticle 1 | **-1.74** | 3.20E-07 | **2.40E-04** |
| **Sfrp5** | secreted frizzled-related sequence protein 5 | **-3.93** | 4.80E-07 | **3.23E-04** |
| **Slc5a7** | solute carrier family 5 (choline transporter), member 7 | **-3.80** | 6.19E-07 | **4.03E-04** |

| | | | | |
|---|---|---|---|---|
| **Dusp9** | dual specificity phosphatase 9 | **-2.06** | 6.82E-07 | **4.29E-04** |
| **Peg3** | paternally expressed 3 | **-2.50** | 7.51E-07 | **4.58E-04** |
| **Zfp618** | zinc finger protein 618 | **-2.74** | 1.25E-06 | **6.99E-04** |
| **Xkr4** | X-linked Kx blood group related 4 | **-2.56** | 1.33E-06 | **7.08E-04** |
| **En1** | engrailed 1 | **-1.42** | 2.11E-06 | **1.00E-03** |
| **Ptch2** | patched 2 | **-2.31** | 3.32E-06 | **1.44E-03** |
| **Mycbp2** | MYC binding protein 2, E3 ubiquitin protein ligase | **-1.35** | 3.57E-06 | **1.45E-03** |
| **Gm49909** | predicted gene, 49909 | **-1.44** | 3.45E-06 | **1.45E-03** |
| **Tuba1a** | tubulin, alpha 1A | **-2.84** | 5.44E-06 | **2.09E-03** |
| **Trim16** | tripartite motif-containing 16 | **-2.08** | 9.07E-06 | **3.05E-03** |
| **Cfap100** | cilia and flagella associated protein 100 | **-1.94** | 1.21E-05 | **3.82E-03** |
| **Tsks** | testis-specific serine kinase substrate | **-2.19** | 1.33E-05 | **4.07E-03** |
| **Gabrr2** | gamma-aminobutyric acid type A receptor subunit rho 2 | **-2.34** | 1.31E-05 | **4.07E-03** |
| **Sez6l** | seizure related 6 homolog like | **-3.63** | 1.45E-05 | **4.35E-03** |
| **Slc1a4** | solute carrier family 1 (glutamate/neutral amino acid transporter), member 4 | **-1.67** | 1.79E-05 | **5.23E-03** |
| **Kcnh2** | potassium voltage-gated channel, subfamily H (eag-related), member 2 | **-2.71** | 1.81E-05 | **5.23E-03** |
| **Sema5b** | sema domain, seven thrombospondin repeats (type 1 and type 1-like), transmembrane domain (TM) and short cytoplasmic domain, (semaphorin) 5B | **-1.61** | 2.09E-05 | **5.67E-03** |
| **Pla2g5** | phospholipase A2, group V | **-2.42** | 2.48E-05 | **6.53E-03** |
| **Cacnb4** | calcium channel, voltage-dependent, beta 4 subunit | **-0.97** | 2.68E-05 | **6.78E-03** |
| **Dennd2b** | DENN domain containing 2B | **-0.76** | 2.93E-05 | **7.27E-03** |
| **Zbtb7c** | zinc finger and BTB domain containing 7C | **-1.20** | 3.00E-05 | **7.32E-03** |
| **Gm3752** | predicted gene 3752 | **-1.15** | 3.42E-05 | **7.89E-03** |
| **Cdon** | cell adhesion molecule-related/down-regulated by oncogenes | **-1.27** | 3.43E-05 | **7.89E-03** |
| **Fam20c** | FAM20C, golgi associated secretory pathway kinase | **-1.84** | 3.46E-05 | **7.89E-03** |
| **Ccdc9b** | coiled-coil domain containing 9B | **-1.91** | 3.80E-05 | **8.43E-03** |
| **D830030K20Rik** | RIKEN cDNA D830030K20 gene | **-1.20** | 5.30E-05 | **1.14E-02** |
| **Fstl3** | follistatin-like 3 | **-1.88** | 5.27E-05 | **1.14E-02** |
| **Gm8895** | predicted gene 8895 | **-3.52** | 6.39E-05 | **1.30E-02** |
| **Vash1** | vasohibin 1 | **-1.01** | 7.96E-05 | **1.52E-02** |
| **Serpine1** | serine (or cysteine) peptidase inhibitor, clade E, member 1 | **-1.81** | 8.61E-05 | **1.57E-02** |
| **Lep** | leptin | **-2.43** | 9.31E-05 | **1.67E-02** |
| **Erc2** | ELKS/RAB6-interacting/CAST family member 2 | **-1.60** | 1.07E-04 | **1.87E-02** |
| **Mageb18** | MAGE family member B18 | **-4.24** | 1.19E-04 | **2.00E-02** |
| **Adam3** | ADAM metallopeptidase domain 3 | **-1.83** | 1.59E-04 | **2.52E-02** |
| **Dclk3** | doublecortin-like kinase 3 | **-2.15** | 1.62E-04 | **2.54E-02** |
| **Gm37240** | predicted gene, 37240 | **-1.06** | 1.77E-04 | **2.64E-02** |
| **Lipf** | lipase, gastric | **-4.25** | 1.75E-04 | **2.64E-02** |
| **Ldlrad4** | low density lipoprotein receptor class A domain containing 4 | **-0.93** | 1.80E-04 | **2.65E-02** |
| **Pcdh20** | protocadherin 20 | **-2.30** | 1.80E-04 | **2.65E-02** |
| **Apbb1** | amyloid beta precursor protein binding family B member 1 | **-1.10** | 1.85E-04 | **2.66E-02** |
| **Rtn4rl1** | reticulon 4 receptor-like 1 | **-1.51** | 1.92E-04 | **2.73E-02** |
| **Pcdh12** | protocadherin 12 | **-1.23** | 2.54E-04 | **3.47E-02** |
| **Gm5941** | predicted gene 5941 | **-4.45** | 2.60E-04 | **3.50E-02** |
| **Synpo2** | synaptopodin 2 | **-1.93** | 2.76E-04 | **3.66E-02** |
| **Mturn** | maturin, neural progenitor differentiation regulator homolog (Xenopus) | **-1.98** | 2.86E-04 | **3.76E-02** |
| **Lama3** | laminin, alpha 3 | **-1.62** | 2.97E-04 | **3.84E-02** |
| **Gli2** | GLI-Kruppel family member GLI2 | **-1.32** | 3.38E-04 | **4.18E-02** |
| **Gm10340** | predicted gene 10340 | **-1.09** | 3.66E-04 | **4.45E-02** |
| **Hr** | lysine demethylase and nuclear receptor corepressor | **-2.29** | 3.67E-04 | **4.45E-02** |
| **Cdc42ep5** | CDC42 effector protein 5 | **-1.42** | 3.70E-04 | **4.45E-02** |
| **Pabpc4l** | poly(A) binding protein, cytoplasmic 4-like | **-1.68** | 3.86E-04 | **4.62E-02** |
| **Rap1gap2** | RAP1 GTPase activating protein 2 | **-1.15** | 4.03E-04 | **4.74E-02** |
| **Oxtr** | oxytocin receptor | **-2.24** | 4.30E-04 | **4.90E-02** |

**Supplementary Table 3: Differentially expressed genes in FLASH-RT mice iscWAT compared to CNT.** Genes with an absolute fold change >1.5 and a false discovery rate (FDR)-adjusted p-value<0.05 were considered differentially expressed.

| Gene Name | Description | log2(FC) | pvalue | padj |
|---|---|---|---|---|
| **Gm20708** | predicted gene 20708 | **6.75** | 1.18E-06 | **1.53E-02** |
| **Vat1l** | vesicle amine transport protein 1 like | **-1.77** | 8.64E-07 | **1.53E-02** |
| **En1** | engrailed 1 | **-1.35** | 3.51E-06 | **3.04E-02** |
